# ChIP-seq analysis of SMAD3 target genes in a sea anemone reveals symbiosis-dependent regulation through TGF-β signaling

**DOI:** 10.64898/2026.11.18.767637

**Authors:** Olivia L. Burleigh, Mascha F. Dix, Maria Ruggeri, Guoxin Cui, Manuel Aranda, Virginia M. Weis

**Affiliations:** Department of Integrative Biology, Oregon State University, Corvallis, OR, United States; Biomedical Sciences Division, Red Sea Research Center, King Abdullah University of Science and Technology, Thuwal, Saudi Arabia; Biological and Environmental Sciences and Engineering Division, Red Sea Research Center, King Abdullah University of Science and Technology, Thuwal, Saudi Arabia

**Keywords:** Cnidarian-algal symbiosis, ChIP-seq, TGF-β, SMAD3, *Exaiptasia diaphana*, Gene expression, Transcription factor, Heat stress

## Abstract

**Background:** Cnidarian-algal symbiosis underpins coral reefs, but elevated temperatures can disrupt it, causing bleaching. Bleaching activates host stress and immune pathways and disrupts symbiotic signaling and nutrient transfer. Among these pathways, the immunosuppressive cytokine transforming growth factor-β (TGF-β) promotes host tolerance of symbionts in cnidarians, but the genes regulated by its transcription factor, SMAD3, remain unknown. SMAD3 ChIP-seq and RNA-seq were used to identify putative targets in symbiotic and aposymbiotic *Exaiptasia diaphana* after 6 and 18 h of heat stress.

**Results:** SMAD3 binding was highly specific to each combination of symbiotic state and heat treatment. Symbiotic state, not heat stress, drove most differential binding, with binding nearly always higher in symbiotic animals. Integration identified 625 putative targets differentially bound and expressed between symbiotic states. Differential binding between symbiotic states peaked at 18 h, when these targets made up the largest share of symbiosis-associated expression differences. Activation targets suggest SMAD3 supports symbiosis through the mucociliary surface, host control of nitrogen available to symbionts, uptake of symbiont-derived retinoids, and immunity. Repression targets, which outnumbered activation targets at 18 h, suggest SMAD3 limits canonical sterol transport, alters G protein-coupled receptor expression, and may reinforce its own activity by repressing negative regulators of TGF-β signaling.

**Conclusions:** SMAD3 appears to act as a bifunctional regulator whose binding is shaped primarily by symbiotic state. Its putative targets suggest roles in cnidarian symbiosis beyond immune suppression, including nutrient exchange and retinoid signaling. These findings provide a framework for TGF-β function in cnidarian symbiosis and thermal stress.

## Background

One of the most significant ecological forms of endosymbiosis is between cnidarians and dinoflagellate algae in the family Symbiodiniaceae [1]. This symbiosis underlies the productivity and calcification that allow reef-building corals to construct the three-dimensional habitat on which the broader reef community depends [2, 3]. During a healthy symbiosis, cnidarian host cells and symbionts exchange inorganic nutrients and photosynthetic products to their mutual benefit. The tight regulation of nutrient recycling between the host and symbiont allows reef-building corals and other cnidarians to thrive in the oligotrophic water of the tropics [4]. A variety of environmental stressors, most notably prolonged elevated temperatures due to climate change, can cause the breakdown of the symbiosis, where host cnidarians lose their algal symbionts in a process known as coral bleaching [5–8]. Extended periods of bleaching without the re-establishment of the symbiosis can lead to disease susceptibility and death, eventually causing reef degradation [6, 9, 10].

The cellular processes underlying bleaching are complex, involving the activation of stress and immune pathways alongside the disruption of symbiotic signaling and nutrient transfer [11]. Under heat stress, the symbiosis breaks down as nutrient exchange between host and symbiont shifts away from its homeostatic balance, destabilizing the nitrogen and carbon cycling that normally sustains the partnership [12]. In addition, heat stress and high light can damage the algae, resulting in the production of reactive oxygen species (ROS) that move into the host, damaging lipids, proteins, and DNA [13, 14]. This damage can trigger apoptosis, necrosis, and autophagy that contribute to symbiont loss [11, 15, 16]. Understanding the full cellular complexity of this symbiotic relationship, both at baseline and at elevated temperatures, is especially pressing given the accelerating rate and severity of global bleaching events driving reef decline in recent years [10].

Host immune suppression to accommodate intracellular symbionts is a recurring theme across symbiotic relationships, reflecting a broadly conserved strategy for establishing and maintaining these partnerships [17, 18]. Previous research has demonstrated the central role of host innate immunity in the regulation of cnidarian-algal symbiosis [19–21]. There is evidence that the cnidarian host immune system is locally suppressed in symbiosis-competent gastrodermal cells during the establishment of symbiosis and this immunosuppression is maintained during stable symbiosis through a variety of cellular pathways, including the downregulation of the master immune regulator transcription factor NF-κB [19, 21–25]. In addition to suppressing pro-inflammatory signaling, host cells promote immune tolerance signaling to allow for symbiont persistence. Transforming growth factor beta (TGF-β) is conserved across metazoans and is a context-dependent tolerogenic cytokine that regulates a variety of cellular processes, including cell differentiation, tissue homeostasis, apoptosis, migration, adhesion, wound healing, and immunity [26–30]. Much of what is known about TGF-β signaling comes from vertebrate systems, where it has been extensively studied for its role in cancer, inflammation, autoimmune disease, and immune suppression [28, 31]. By comparison, its role in symbiotic cnidarians has received far less attention, though existing studies implicate TGF-β signaling in regulating symbiosis and immunity [32–34].

In the sea anemone *Exaiptasia diaphana* (Aiptasia), blocking TGF-β signaling with an anti-TGF-β antibody increased immune stimulation, as evidenced by elevated nitric oxide production [33]. In addition, antibody incubations prevented symbionts from successfully proliferating in aposymbiotic Aiptasia, indicating that tolerogenic TGF-β activity is required for symbiont establishment. Conversely, adding exogenous TGF-β partially reduced the bleaching response in heat-stressed, symbiotic anemones, suggesting that TGF-β-mediated immune tolerance also helped maintain symbiosis under thermal stress. Fuess et al. [34] found that enhancing TGF-β signaling in the coral *Orbicella faveolata* suppressed the host’s immune transcriptional response to bacterial challenge, while inhibiting TGF-β preserved and even strengthened it, revealing a trade-off between TGF-β-mediated tolerance and immune defense. These findings support TGF-β as a conserved immunosuppressive regulator in corals, and with the complementing evidence from Aiptasia, link the pathway to both symbiosis and thermal tolerance across cnidarians.

TGF-β signaling is propagated through SMAD proteins, the canonical intracellular transducers of the pathway (Fig. 1). In vertebrates, the TGF-β ligand binds to TGF-β serine/threonine kinase receptors which in turn phosphorylate SMAD3. Phosphorylated SMAD3 then partners with SMAD4, and this complex translocates to the nucleus to regulate transcription alongside coactivators, corepressors, and other transcription factors bound at target loci [29, 30]. Most TGF-β pathway components, including SMAD proteins, have been identified in the Aiptasia genome [33, 35]. Although the TGF-β ligand has been linked to symbiosis, immunity, and thermal tolerance in cnidarians, it does not act on target genes directly; its downstream effects are executed by the SMAD complex and the specific genes the complex binds. Most cnidarian studies to date have measured pathway activity through ligand-level manipulation alone, leaving the actual transcriptional mechanism undefined. It is still unknown which genes SMAD3 binds, under which physiological conditions, and how that binding shifts between symbiotic states and under thermal stress.

**Figure 1.**
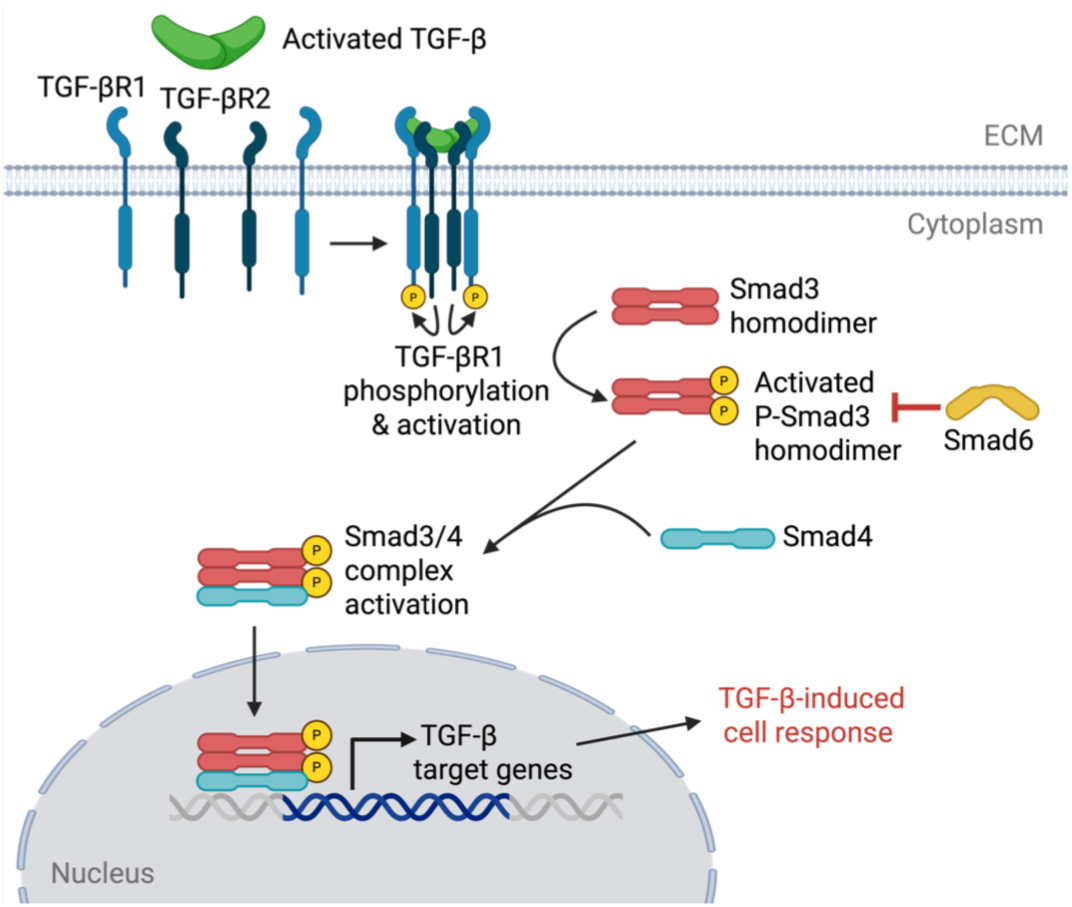
TGF-β/SMAD signaling pathway in Aiptasia based on genomic searches. Activated TGF-β ligand binds the type II receptor (TGF-βR2), which recruits the type I receptor (TGF-βR1) and activates it through phosphorylation. Activated TGF-β receptors then phosphorylate SMAD3. The inhibitory SMAD6 can negatively regulate the pathway by inhibiting SMAD3 phosphorylation. Phosphorylated SMAD3 (P-SMAD3) forms a complex with the common mediator SMAD4. The activated SMAD3/4 complex translocates to the nucleus and regulates transcription of TGF-β target genes, producing a TGF-β-induced cellular response. The polyclonal antibody used for ChIP-seq was raised against Aiptasia SMAD3 and recognizes SMAD3 within this complex.

Advances in high-throughput sequencing have opened new avenues for probing these questions directly in cnidarian systems, with genomic and transcriptomic approaches substantially advancing understanding of the molecular basis of cnidarian-algal symbiosis [36]. Chromatin immunoprecipitation sequencing (ChIP-seq) offers a direct means of mapping genome-wide transcription factor-DNA interactions at a resolution unavailable through pathway manipulation studies alone. Here we paired SMAD3 ChIP-seq with RNA-seq to characterize SMAD3 binding and identify its downstream transcriptional targets across symbiotic states and thermal treatments in Aiptasia, with the aim of understanding how TGF-β/SMAD3 signaling contributes to the maintenance and thermal sensitivity of cnidarian-algal symbiosis.

## Results

### SMAD3 binds to distinct genomic regions with minimal overlap between conditions

Conditions are abbreviated by treatment (C, control; H, heat), symbiotic state (A, aposymbiotic; S, symbiotic), and duration of heat stress in hours (6 or 18) (Fig. 2A). SMAD3 binding was greatest in CS with 5,161 consensus peaks and CA with 4,789 peaks (Fig. 3A). HS6 and HS18 showed a pronounced reduction in reproducible binding sites with 1,626 and 1,394 peaks, respectively, while HA6 and HA18 declined more modestly with 4,212 and 3,842 peaks, respectively (Fig. 3A). Despite these differences in total peak number, 86% of consensus peaks were unique to a single condition (Fig. 3A). Only 13 peaks were shared across all six conditions. Shared peaks more often grouped by symbiotic state than by treatment: 49 peaks were found only in all three symbiotic conditions and 34 only in all three aposymbiotic conditions, whereas a single peak was shared exclusively among the four heat-treated conditions. The two control conditions shared 208 peaks exclusively. This pattern was consistent under a stricter reproducibility threshold requiring all three replicates (c = 3; 93% condition-unique; Additional file 1: Fig. S3).

**Figure 2.**
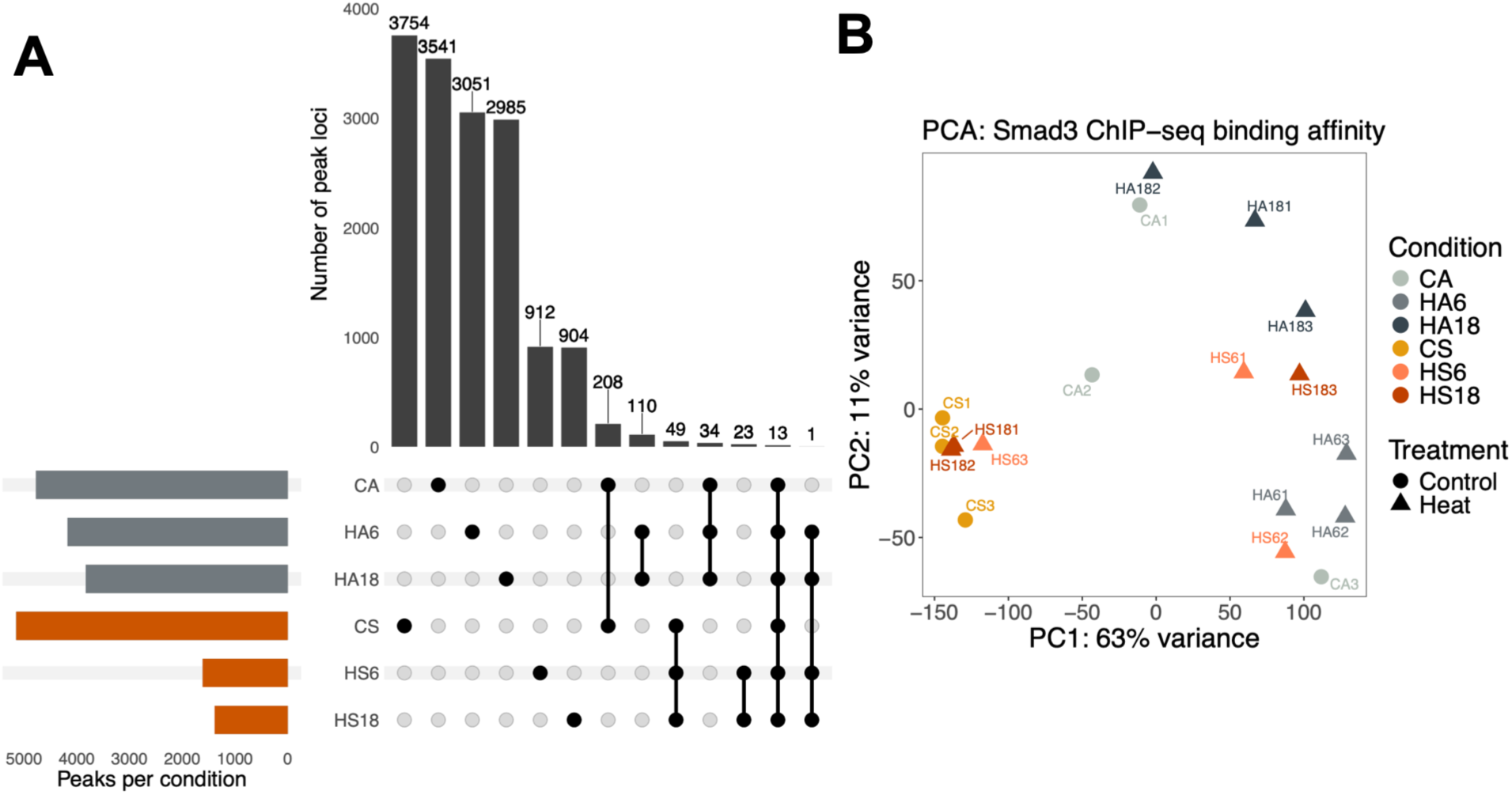
Experimental design and ChIP-seq sampling methods. (**A**) The temperature ramp from 25°C to 34°C, depicted as a black line. Heat-stressed animals were sampled at 34°C after 6 h or 18 h (red arrows); controls were held at 25°C and sampled at the 6 h timepoint only (blue arrow). Condition abbreviations indicate treatment (C, control; H, heat), symbiotic state (A, aposymbiotic; S, symbiotic), and hours of heat stress (6 or 18): CA, CS, HA6, HS6, HA18, and HS18 (colored boxes; n = 3 per condition). (**B**) ChIP-seq method diagram showing immunocomplex recovery of SMAD3 protein complex and library preparation for SMAD3 and input libraries.

**Figure 3.**
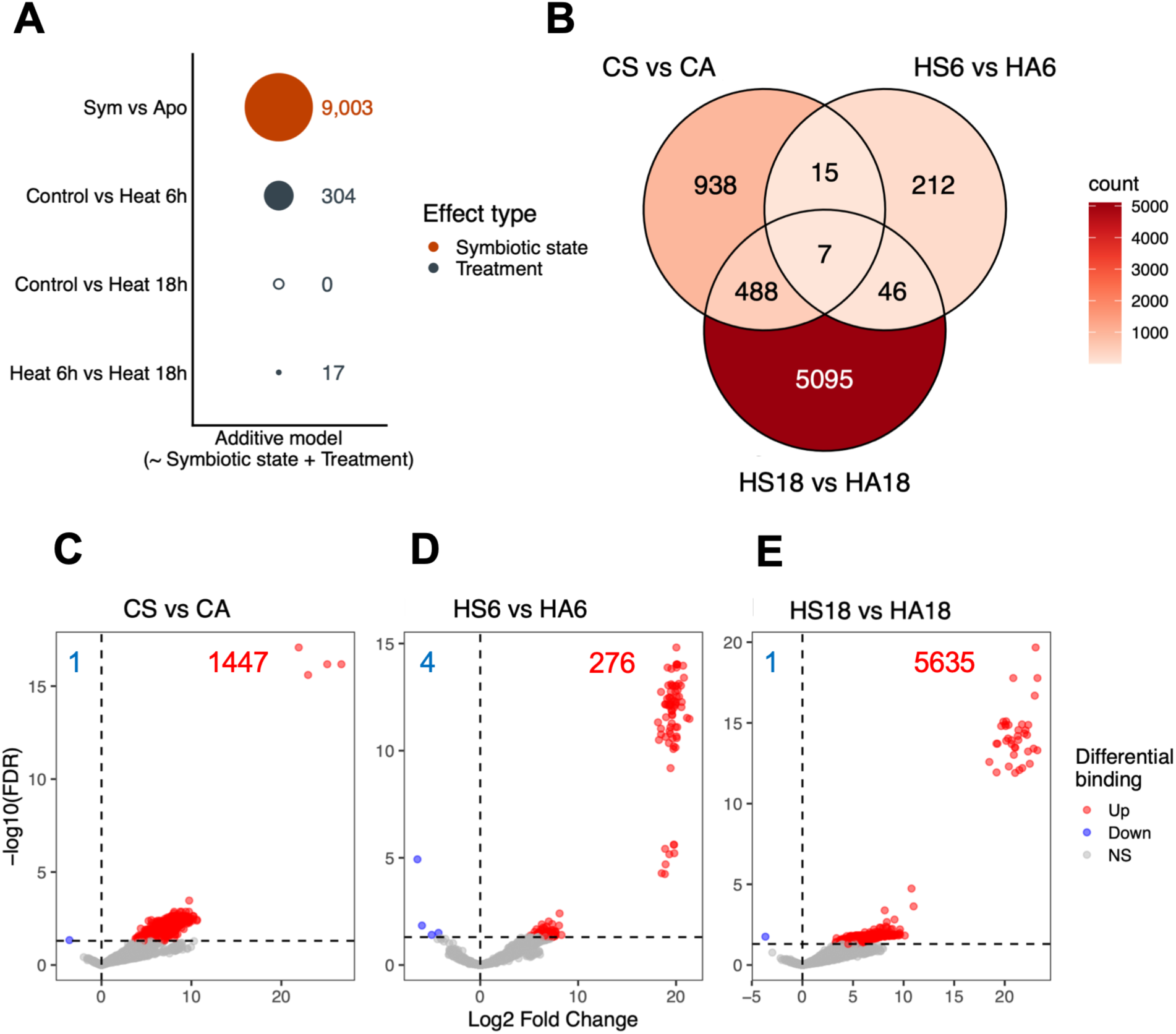
SMAD3 binds to distinct genomic regions. (**A**) UpSet plot of Multiple Sample Peak Calling (MSPC) consensus peaks by condition, restricted to peaks recovered in two out of three replicates; total consensus peaks per condition are shown at left. Grey, aposymbiotic; orange, symbiotic. Specific intersections between conditions of interest are shown. (**B**) Principal component analysis of SMAD3 ChIP-seq binding affinity from variance-stabilized read counts at consensus peak loci (PC1, 63%; PC2, 11% of variance). Symbiotic state separated by color (grey, orange) and treatment separated by shape.

### Symbiotic state explains most SMAD3 differential binding

PCA of blind variance-stabilized SMAD3 ChIP-seq read counts across the 17,569 consensus binding sites separated samples primarily by symbiotic state along PC1, which explained 63% of the variance (Fig. 3B). PERMANOVA showed that both symbiotic state (R² = 0.22, F = 5.32, p = 0.007) and heat treatment (R² = 0.19, F = 2.24, p = 0.038) explained significant variance in SMAD3 binding, with no significant interaction (R² = 0.096, p = 0.286). Overall dispersion did not differ significantly by symbiotic state or treatment (betadisper, p = 0.477 and p = 0.691, respectively), although three heat-stressed symbiotic samples, HS6-1, HS6-2, and HS18-3, appeared as outliers in ordination space, overlapping with the aposymbiotic cluster.

Differential binding analysis distinguished the effects of symbiotic state and heat more clearly. Under an additive model (design = ∼ symbiotic state + treatment), symbiotic state accounted for 9,003 differentially bound peaks, compared with at most 304 for any heat treatment comparison (Fig. 4A). Because the additive model assumes a single symbiotic-state effect shared across treatments, an interaction model (design = ∼ symbiotic state × treatment) was also fit to compare symbiotic states at each timepoint and heat treatments within each symbiotic state. Heat treatment alone produced few differentially bound peaks within either symbiotic state (Additional file 1: Fig. S4). In aposymbiotic animals, the contrast HA6 vs CA had 90 differentially bound peaks, HA18 vs CA had 102 peaks, and HA6 vs HA18 had 480 peaks (Additional file 1: Fig. S4A–C). In symbiotic animals, there were 63 differentially bound peaks between HS6 and CS, all of which were higher in CS (Additional file 1: Fig. S4D). There were no peaks differentially bound between HS18 and CS or between HS6 and HS18 (Additional file 1: Fig. S4E–F). Only a small subset of peaks had a significant symbiotic state × treatment interaction, with 97 peaks at 6 h and 41 peaks at 18 h.

**Figure 4.**
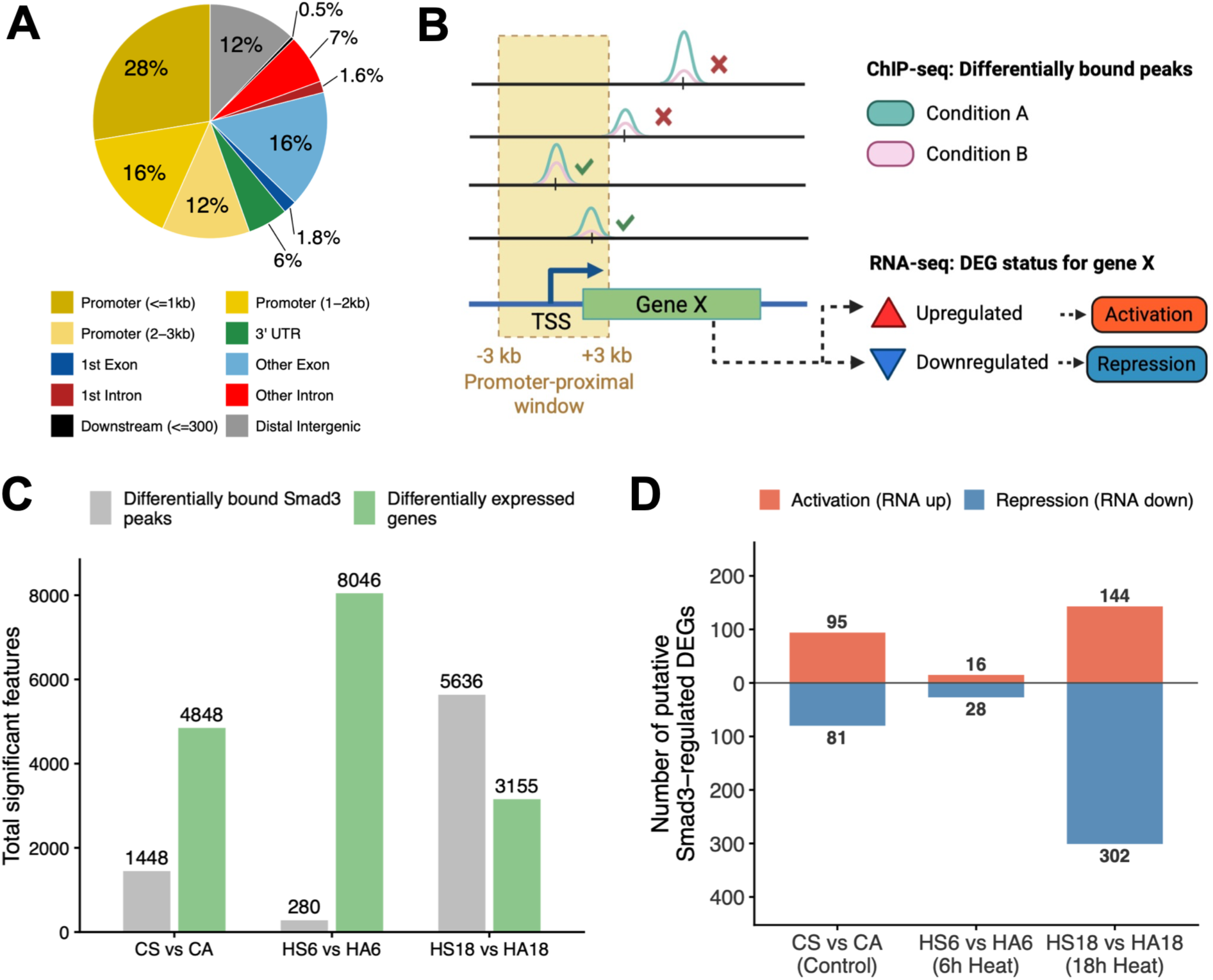
Differential SMAD3 binding is driven by symbiotic state and extends to more sites under prolonged heat stress. (**A**) Number of differentially bound SMAD3 peaks (FDR < 0.05) from the additive model (design = ∼symbiotic state + treatment) in DiffBind. Bubble size scales with peak count. Orange, symbiotic state effect; grey, heat treatment effect. Open circles indicate zero significant peaks. (**B**) Venn diagram of differentially bound peaks from the interaction model (design = ∼symbiotic state × treatment) across the three symbiotic state contrasts (Sym vs Apo at each treatment level); fill color scales with peak count. (**C–E**) Volcano plots of differential SMAD3 binding for each symbiotic state contrast from the interaction model. X-axis, log2 fold change (Sym relative to Apo); y-axis, −log10(FDR). Red, increased binding in symbiotic animals; blue, decreased binding; grey, not significant (NS). Numbers indicate the count of significant peaks per direction. The dashed horizontal line indicates the significance threshold FDR = 0.05.

Contrasts between symbiotic states at each timepoint yielded more differentially bound peaks than most heat-specific contrasts within a symbiotic state. The CS vs CA and HS18 vs HA18 contrasts shared the most peaks: 495 of the 1,448 peaks differentially bound under control conditions (34%) were also differentially bound at 18 h, although these made up only 9% of the 5,636 peaks in the HS18 vs HA18 contrast, which were otherwise largely unique to 18 h (Fig. 4B). In contrast, the smaller HS6 vs HA6 contrast was largely distinct, with 212 of 280 peaks (76%) unique to 6 h, 53 shared with the 18 h contrast, and 22 shared with the control contrast (Fig. 4B). Only 7 peaks were differentially bound across all three symbiotic-state contrasts. Together, these results indicate that a subset of symbiotic-state differences under control conditions persisted at 18 h, while most differentially bound peaks at each timepoint were specific to that timepoint.

Nearly all differentially bound peaks across symbiotic-state contrasts had higher SMAD3 binding in symbiotic compared to symbiotic animals (Fig. 4C–E). Binding was higher in symbiotic animals at 1,447 of 1,448 peaks in the CS vs CA contrast, 276 of 280 peaks in the HS6 vs HA6 contrast, and 5,635 of 5,636 peaks in the HS18 vs HA18 contrast. The number of differentially bound peaks was greatest after 18 h of heat stress, indicating that higher SMAD3 binding in symbiotic animals extends to more sites under prolonged heat.

### SMAD3 is associated with both activation and repression of target genes

Over half of SMAD3 consensus peaks, 55.4%, were located within promoter-proximal regions defined as ±3 kb of the TSS (Fig. 5A). The highest concentration fell within 1 kb of the TSS (28%), followed by 1–2 kb (16%), and 2–3 kb (12%). The remaining peaks were distributed across exon regions (18.0%), distal intergenic regions (12%), intron regions (8.6%), and UTRs (5.6%). The frequent binding of SMAD3 near promoters is consistent with its role in directly regulating gene expression at promoter-proximal regulatory elements, where SMAD3 recruits chromatin-modifying cofactors to activate or repress TGF-β-responsive genes [37].

**Figure 5.**
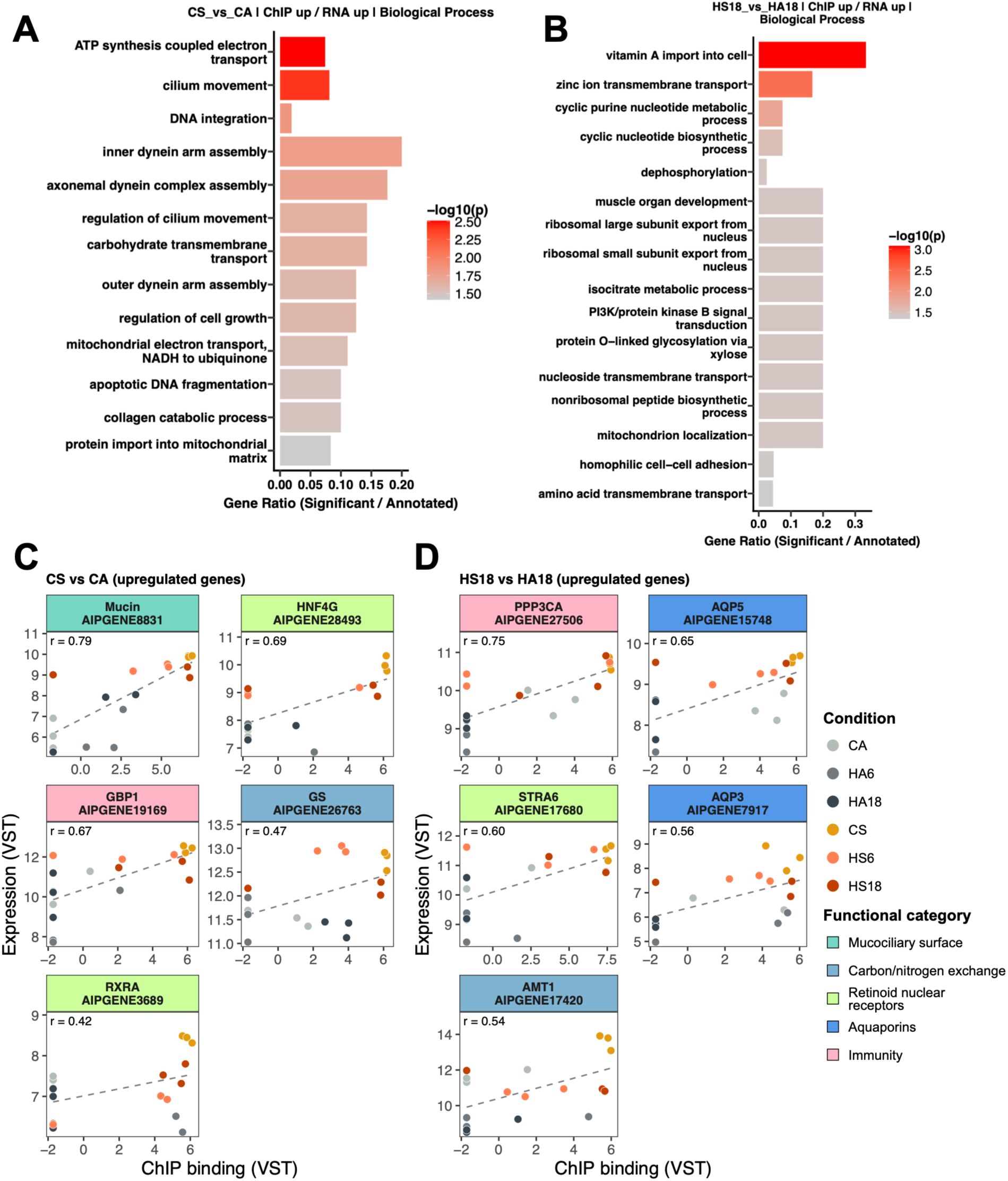
Integration of SMAD3 binding and gene expression. (**A**) Genomic annotation of Multiple Sample Peak Calling (MSPC) consensus peaks across all conditions (ChIPseeker), with 55.4% falling within 3 kb of a transcription start site. 5′ UTR is omitted (0.03%). (**B**) ChIP-seq/RNA-seq integration workflow. Differentially bound SMAD3 peaks whose midpoint fell within ±3 kb of a TSS were assigned to the corresponding gene; peaks outside this promoter-proximal window were excluded. Peak-associated genes were cross-referenced with RNA-seq differential expression for the same contrast. Genes with increased SMAD3 binding and upregulated expression were classified as putative activation targets and those with increased binding and downregulated expression were classified as putative repression targets. (**C**) Number of significant features per symbiotic state contrast. Grey, differentially bound SMAD3 peaks; green, DEGs used in ChIP-seq/RNA-seq integration analysis. (**D**) Number of putative SMAD3-targeted DEGs per contrast. Contrasts compare symbiotic vs aposymbiotic animals at each timepoint (control, 6 h heat, 18 h heat). Bars show the number of DEGs carrying a differentially bound SMAD3 peak in their promoter-proximal region (±3 kb of the TSS) for the same contrast. Red, activation (gene expression up, binding up); blue, repression (gene expression down, binding up).

Condition contrasts between symbiotic states were prioritized for integrating the ChIP-seq and RNA-seq datasets, as these contrasts yielded the largest numbers of differentially bound peaks and thus had the greatest power to detect concordance between SMAD3 binding and gene expression. Within each contrast, differentially bound peaks (padj < 0.05) were restricted to those whose midpoint fell within a promoter-proximal window, defined as ±3 kb of a TSS. Promoter proximity was used to prioritize peaks most likely to influence a nearby gene, but it does not establish direct regulation or identify a unique target gene, so peak–gene pairs were treated as putative associations (Fig. 5B). This retained 653, 113, and 2,642 promoter-proximal peaks in CS vs CA, HS6 vs HA6, and HS18 vs HA18 contrasts, respectively (Additional file 3: Table S2B). When a peak fell within the promoter-proximal window of more than one gene, it was assigned to each gene, as this analysis cannot distinguish which gene the peak is more likely to regulate. As a result, the number of associated genes exceeded the number of promoter-proximal peaks, with 864, 135, and 3,260 unique genes in the three contrasts, respectively (Additional file 3: Table S2B). Genes carrying a differentially bound promoter-proximal SMAD3 peak were then cross-referenced with DEGs from the matched RNA-seq contrast (see Additional file 1: Fig. S5–10 for RNA-seq results).

Because the ChIP-seq and RNA-seq pipelines use different gene identifiers, integration was limited to DEGs that could be matched to the gene models used for peak annotation. This reduced the 7,876, 12,319, and 5,535 total DEGs to 4,848, 8,046, and 3,155 matched DEGs in the three contrasts, respectively (Additional file 3: Table S2A–B). Peaks with decreased SMAD3 binding in symbiotic animals were rare (Fig. 4C–E), and none fell within the promoter-proximal window of a matched DEG, so no integrated genes were associated with reduced binding due to symbiosis. All integrated genes therefore carried a peak with greater SMAD3 binding in symbiotic animals and were classified as putative activation targets when the gene’s expression was upregulated and putative repression targets when downregulated (Fig. 5B).

In the CS vs CA contrast, intersecting the 864 promoter-associated genes with the 4,848 matched DEGs returned 176 putative SMAD3-targeted DEGs, comprising 95 activation and 81 repression targets (Fig. 5C–D; Additional file 3: Table S2B). The same intersection returned 44 targets in the HS6 vs HA6 contrast (135 promoter-associated genes and 8,046 matched DEGs; 16 activation and 28 repression) and 446 targets in the HS18 vs HA18 contrast (3,260 and 3,155; 144 activation and 302 repression) (Fig. 5C–D). Across all three contrasts, these putative targets shifted toward repression under heat, reaching approximately twice as many repression targets at 18 h (Fig. 5D). The proportion of matched DEGs that were putative SMAD3 targets was lowest at 6 h (0.5%) and highest at 18 h (14.1%), compared with 3.6% at control (Additional file 3: Table S2B). Across the three symbiotic-state contrasts, 625 unique genes were identified as putative SMAD3 targets. Of these, 587 genes were targets in only one contrast, 35 were targets in two contrasts, and three were targets in all three contrasts, giving 666 targets in total when each contrast is counted separately (Additional file 7). Assigning peaks directly to AIPGENE gene models without identifier matching identified 1,140 gene-contrast pairs, recovering 521 of the 666 from the main analysis with the same regulatory mode. The additional targets were mostly uncharacterized repression targets, indicating that the main analysis is likely underestimates repression targets (Additional file 3: Table S2C).

GO enrichment of SMAD3 target genes was most informative for the contrasts CS vs CA and HS18 vs HA18, as the remaining contrasts did not yield enough significant genes for robust GO enrichment. Enrichment was further constrained by the annotation itself: only 64.2% of genes in the Aiptasia genome carry associated GO terms, so the target list for each contrast was restricted to GO-annotated genes only, reducing the counts below those reported in the preceding section. Additionally, of the SMAD3 target DEGs (Fig. 5D), 49.6% were uncharacterized and not informative for GO enrichment.

To describe how consistently binding and expression co-varied across individual samples, Pearson correlation was calculated between variance-stabilized counts of SMAD3 ChIP-seq binding and transcript expression across all 18 samples for every target gene in the symbiotic state contrasts (Additional file 7). A select set of genes with some of the highest correlation values is displayed in Figs. 6C–D and 7C-D. Because samples were grouped by condition, these correlations largely reflect the difference between symbiotic states rather than variation within conditions.

**Figure 6.**
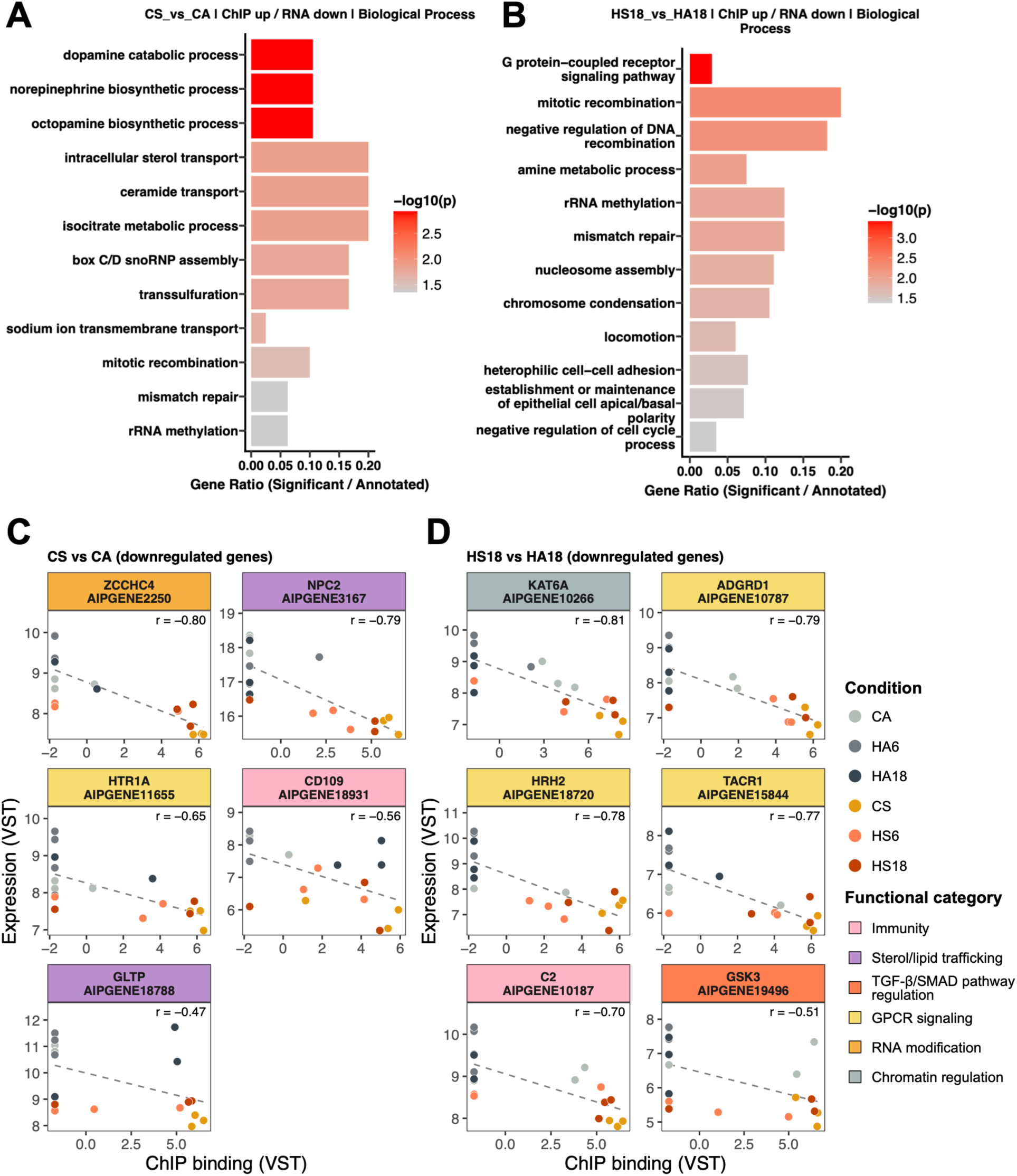
Putative SMAD3 activation targets in symbiotic animals. (**A–B**) Biological process GO terms enriched using topGO for genes that contain a differentially bound SMAD3 peak in their promoter-proximal region and are upregulated in symbiotic animals in the contrasts CS vs CA (**A**) or HS18 vs HA18 (**B**), respectively. Bar color indicates significance (−log10 weight01 Fisher p-value) with a significance threshold of p < 0.05; bar length shows the gene ratio (significant genes/annotated genes per GO term). (**C–D**) Pearson correlation between SMAD3 ChIP-seq binding intensity (x-axis) and transcript expression (y-axis) for select putative activation target genes in the CS vs CA (**C**) and HS18 vs HA18 (**D**) contrasts. Both axes display variance-stabilized transformed counts. Each point represents one of 18 samples, colored by condition (aposymbiotic in grey, symbiotic in orange). Pearson r is shown in parentheses for each gene. The dashed line indicates the linear fit. Panel background colors group genes by functional category: teal, mucociliary surface; light blue, carbon/nitrogen exchange; green, retinoid nuclear receptors; dark blue, aquaporins; pink, immunity.

**Figure 7.**
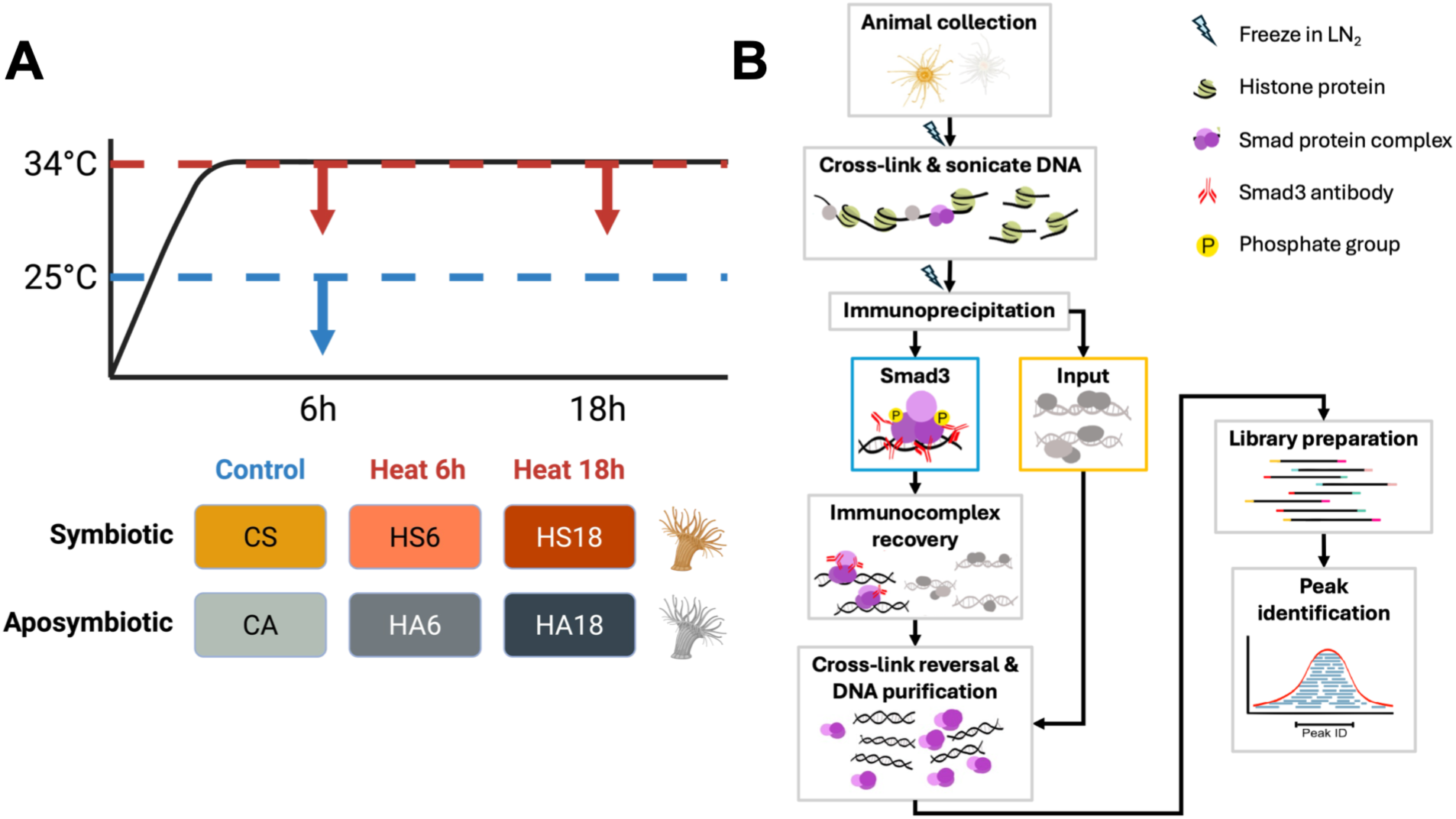
Putative SMAD3 repression targets in symbiotic animals. (**A–B**) Biological process GO terms enriched using topGO for genes that contain a differentially bound SMAD3 peak in its promoter-proximal region and are downregulated in symbiotic animals in the contrasts CS vs CA (**A**) or HS18 vs HA18 (**B**), respectively. Bar color indicates significance (−log10 weight01 Fisher p-value); x-axis shows the gene ratio (significant genes/annotated genes per GO term). (**C–D**) Pearson correlation between SMAD3 ChIP-seq binding intensity (x-axis) and transcript expression (y-axis) for select putative repression target genes in the CS vs CA (**C**) and HS18 vs HA18 (**D**) contrasts. Both axes display variance-stabilized transformed counts. Each point represents one of 18 samples, colored by condition (aposymbiotic in grey, symbiotic in orange tones). Pearson r is shown in parentheses for each gene. The dashed line indicates the linear fit. Panel background colors group genes by functional category: pink, immunity; purple, sterol/lipid trafficking; red-orange, TGF-β/SMAD pathway regulation; yellow, GPCR signaling; orange, RNA modification; grey, chromatin regulation.

### SMAD3 binding is associated with upregulation of genes in symbiotic animals

Out of the 57 informative DEGs with elevated SMAD3 binding and upregulated expression in the CS vs CA contrast, the most enriched biological process terms included ATP synthesis-coupled electron transport (GO:0042773) and ciliary movement (GO:0003341) (weight01 Fisher p < 0.01) with 13 significant terms total (Fig. 6A). Out of the 95 informative DEGs with elevated SMAD3 binding and upregulated expression in the HS18 vs HA18 contrast, the most enriched biological process terms included vitamin A import into cell (GO:0071939) and zinc ion transmembrane transport (GO:0071577) (weight01 Fisher p < 0.01) with 16 significant terms total (Fig. 6B).

Putative activation targets spanned functional categories including mucociliary surfaces, carbon/nitrogen exchange, retinoid nuclear receptors, aquaporins, and immunity, among others. In the CS vs CA contrast (Fig. 6C), genes with positive Pearson correlations include mucin-like protein (AIPGENE8831; r = 0.79), guanylate-binding protein 1 (GBP1; AIPGENE19169; r = 0.67), hepatocyte nuclear factor 4-gamma (HNF4γ; AIPGENE28493; r = 0.69), retinoid X receptor RXR-alpha-A (RXRα; AIPGENE3689; r = 0.42), and glutamine synthetase (GS; AIPGENE26763; r = 0.47). In the HS18 vs HA18 contrast (Fig. 6D), genes with positive Pearson correlations include serine/threonine-protein phosphatase 2B catalytic subunit alpha isoform (PPP3CA; AIPGENE27506; r = 0.75), aquaporin-5 (AQP5; AIPGENE15748; r = 0.65), aquaporin-3 (AQP3; AIPGENE7917; r = 0.56), stimulated by retinoic acid gene 6 protein-like (STRA6; AIPGENE17680; r = 0.60), and putative ammonium transporter 1 (AMT1; AIPGENE17420; r = 0.54).

### SMAD3 binding is associated with downregulation of genes in symbiotic animals

In contrast to the activation set, genes with elevated SMAD3 binding but reduced expression in symbiotic animals are consistent with a repressive role for SMAD3. Out of the 40 informative DEGs with elevated binding but downregulated expression in the CS vs CA contrast, enriched terms included dopamine catabolic process (GO:0042420), norepinephrine biosynthetic process (GO:0042421), and octopamine biosynthetic process (GO:0006589) (weight01 Fisher p < 0.01) with 12 significant terms total (Fig. 7A). Among the 136 DEGs with elevated binding but downregulated expression in the HS18 vs HA18 contrast, enriched terms include G protein-coupled receptor signaling pathway (GO:0007186), mitotic recombination (GO:0006312), negative regulation of DNA recombination (GO:0045910), and amine metabolic process (GO:0009308) (weight01 Fisher p < 0.01) with 12 significant terms total (Fig. 7B).

Putative repression targets spanned functional categories including immunity, sterol/lipid trafficking, TGF-β pathway regulation, GPCR signaling, RNA modification, and chromatin regulation, among others. In the CS vs CA contrast (Fig. 7C), genes with negative Pearson correlations include rRNA N6-adenosine-methyltransferase ZCCHC4 (ZCCHC4; AIPGENE2250; r = −0.80), canonical NPC intracellular cholesterol transporter 2 (NPC2; AIPGENE3167; r = −0.79), glycolipid transfer protein (GLTP; AIPGENE18788; r = −0.47), 5-hydroxytryptamine receptor 1A (HTR1A; AIPGENE11655; r = −0.65), and CD109 antigen (CD109; AIPGENE18931; r = −0.56). In the HS18 vs HA18 contrast (Fig. 7D), genes with negative Pearson correlations include histone acetyltransferase KAT6A (KAT6A; AIPGENE10266; r = −0.81), adhesion G-protein coupled receptor D1 (ADGRD1; AIPGENE10787; r = −0.79), histamine H2 receptor (HRH2; AIPGENE18720; r = −0.78), substance-P receptor (TACR1; AIPGENE15844; r = −0.77), complement C2 (C2; AIPGENE10187; r = −0.70), and a putative glycogen synthase kinase-3 homolog (GSK3; AIPGENE19496; r = −0.51).

### Functional prediction of SMAD3 binding

To independently assess the putative SMAD3 targets identified by promoter-proximal intersection, BETA [38] was used to infer SMAD3 regulatory function and predict direct targets for each symbiotic-state contrast. SMAD3 binding was significantly associated with both transcriptional activation and repression in all three contrasts, with activation p-values of 0.0085, 0.0116, and 0.0013 and repression p-values of 0.0002, 0.0030, and 0.0165 for the CS vs CA, HS6 vs HA6, and HS18 vs HA18 contrasts, respectively (Additional file 1: Fig. S11). Across the three symbiotic-state contrasts, 63–91% of putative activation and repression targets in each contrast (85% overall) were independently recovered as BETA-predicted direct targets of SMAD3 with the same direction of regulation, including 18 of the 21 candidate genes shown in Fig. 6C–D and 7C–D (Additional file 7).

## Discussion

This study identified putative targets of SMAD-dependent TGF-β signaling in Aiptasia by pairing SMAD3 ChIP-seq with RNA-seq, classifying bound genes that were upregulated or downregulated in symbiotic animals as putative activation or repression targets, respectively. SMAD3 differential binding was highly context-dependent and associated more strongly with symbiotic state than with heat stress. Its binding was associated with both activated and repressed genes, suggesting a bifunctional role of SMAD3. These results suggest how the pleiotropic cytokine TGF-β may support symbiosis and the response to short-term heat stress, while highlighting the complexity of transcription factor dynamics in a whole-animal system.

### SMAD3 binding modulates symbiosis more than the heat stress response

In cnidarians, TGF-β signaling promotes host tolerance of symbionts. In Aiptasia, blocking TGF-β increased immune responses and prevented symbiont colonization, whereas exogenous TGF-β suppressed immune activation [33], and pathway inhibition similarly reduced colonization in coral larvae [32]. Consistent with greater TGF-β activity in symbiotic hosts, symbiotic state separated samples along PC1 and accounted for far more differentially bound peaks than thermal treatment (Fig. 3B; Fig. 4A), and nearly all differentially bound peaks had higher SMAD3 binding in symbiotic animals (Fig. 4C–E). Because most consensus peaks were unique to a single condition (Fig. 3A), symbiosis appears to concentrate SMAD3 binding at a subset of sites rather than increase binding genome-wide.

Less is known about TGF-β signaling during heat stress, although exogenous TGF-β partially reduced bleaching in heat-stressed Aiptasia [33]. In this study, heat produced few differentially bound peaks within either symbiotic state, although variability among heat-stressed symbiotic replicates may have limited detection (Fig. 3B; Additional file 1: Fig. S4). The early heat response instead appeared to involve other pathways. The heat treatment in the present study induced a proteostasis response in both symbiotic states, consistent with activation of HSF1 [39] (Additional file 1: Figs. S8-S9). NF-κB and TNF signaling terms were enriched at both 6 h and 18 h in heat-stressed aposymbiotic animals, whereas in symbiotic animals only a single NF-κB term was enriched at 18 h, consistent with previous work showing lower NF-κB activity in symbiotic Aiptasia [19] (Additional file 1: Figs. S8–S9). In contrast, binding differences between symbiotic states were greatest at 18 h, when SMAD3 targets also accounted for a growing share of symbiosis-associated expression differences, whereas differential expression peaked at 6 h (Fig. 4B; Fig. 5C–D; Table S2). Together with the protective effect of TGF-β against bleaching, this suggests SMAD3 may help sustain symbiotic-state differences during prolonged heat stress rather than mediate the acute heat response.

### Chromatin context and cofactors may explain SMAD3 binding patterns

Differences in chromatin accessibility and cofactor availability may underlie this redistribution of SMAD3 binding across conditions. Chromatin accessibility, histone modifications, and DNA methylation all shape symbiosis and thermal stress responses in cnidarians [40–42], in part by governing where transcription factors can bind [30, 43]. In Aiptasia, accessible regions in symbiotic animals were enriched for SMAD2/3/4 motifs across temperatures [41]. Our results extend this distinct symbiotic-state landscape from accessibility to SMAD3 binding itself.

SMAD proteins recognize short DNA motifs that occur throughout the genome and therefore depend on cofactors for target selection [44–46]. The canonical CAGAC SMAD-binding element was not enriched in our peaks in any condition, including among the most enriched peaks (see Additional file 1 for HOMER methods; Additional file 4). This contrasts with the SMAD motif enrichment reported in symbiotic accessible chromatin [41] and suggests that motif presence alone does not determine where SMAD3 binds. SMAD proteins also bind the GC-rich 5GC motifs GGCGC and GGCCG with affinity comparable to the CAGAC element. These motifs are more prevalent than CAGAC at SMAD-bound regions in mammalian cells and are recognized by SMAD4 from the early-diverging placozoan *Trichoplax adhaerens* [47]. Because 5GC motifs are short, GC-rich, and often clustered, their enrichment may be masked by GC-matched background models, so direct SMAD3 binding may have gone undetected. Motifs for p53 and ETS-family factors, known SMAD2/3 partners in vertebrates [43], showed modest enrichment of no more than 1.4-fold over background, with p53 enriched only in CS and ETS-family motifs enriched in CS and HA18 (Additional file 4). These condition-dependent patterns are consistent with cofactor-mediated recruitment of SMAD3, although the small effect sizes warrant validation. Whole-animal ChIP-seq also averages enrichment signals across cell types, which may dilute both cell-type-specific binding and the motif signatures that accompany it.

### Linking SMAD3 binding to gene expression

Integrating differential binding with differential expression identified 625 unique putative SMAD3-targeted genes across the three symbiotic-state contrasts (Fig. 5D; Additional file 7). These genes represent a small fraction of SMAD3 binding events. Most differentially bound peaks fell outside promoter-proximal windows, and most promoter-associated genes were not differentially expressed (Fig. 5C–D; Additional file 3: Table S2). This is expected, as transcription factors bind far more sites than they regulate, and only a minority of binding events produce measurable changes in the expression of nearby genes [48]. Binding without a detectable expression change may reflect sites that prime genes for later activation, regulation shared with other transcription factors, or effects restricted to certain cell types that are diluted in whole-animal samples. Our integration therefore prioritized the binding events with the strongest regulatory evidence. Two analyses support this prioritized set. Binding and expression of target genes co-varied consistently across individual samples, although this largely reflects the selection of targets for differences between symbiotic states and the grouping of samples by condition. More independently, BETA, which scores all consensus peaks by distance to the TSS rather than relying on differentially bound peaks, recovered 85% of gene-contrast pairs as predicted targets with the same direction of regulation (Additional file 7). Both analyses show concordance rather than causation, so these genes are considered putative SMAD3 targets that require further experimental validation.

### SMAD3 activation targets suggest roles in symbiosis maintenance

SMAD3 activation targets in symbiotic animals point toward symbiosis maintenance rather than an activated stress response. Under control conditions, activated genes in symbiotic animals were enriched for ciliary movement and dynein assembly alongside mitochondrial electron transport, suggesting coordinated investment in motile cilia and the energy that powers them. In sea anemones, ciliated pharyngeal grooves called siphonoglyphs pump seawater into the gastrovascular cavity to inflate the body column and maintain the hydrostatic skeleton [49]. Ciliary currents within this cavity also circulate fluid, food particles, and cells across the gastrodermis that houses symbionts, and in colonial cnidarians transport fixed carbon and symbiont-containing cells between polyps [50, 51]. On the outer surface, mucus forms a protective layer that traps particles and harbors host-associated microbes [52], and ciliary flows move this layer across the epidermis to maintain the surface microbial community and remove pathogens [53]. A mucin-like protein was among the most strongly correlated activation targets under control conditions, suggesting that, together with the ciliary genes, SMAD3 binding activity may contribute to maintaining this mucociliary surface in symbiotic animals.

Retinoid-related genes were among SMAD3 activation targets in both contrasts. Under prolonged heat, activation targets were enriched for vitamin A import into the cell, reflecting activation of STRA6-like retinol transporters, and under control conditions they included the retinoid receptor RXRα. Animals cannot synthesize vitamin A and instead derive it from carotenoids such as β-carotene, which Symbiodiniaceae produce [54, 55]. Activation of these genes in symbiotic animals is consistent with host uptake of symbiont-derived retinoids, although the role of retinoid signaling in cnidarian-dinoflagellate symbiosis remains to be tested.

Other activation targets were involved in nitrogen assimilation, membrane transport, and immunity. GS, which assimilates ammonium into glutamine, was activated under control conditions, and the ammonium transporter AMT1 was activated following 18 h of heat stress. In Aiptasia, symbiont-derived glucose induces both genes to drive host ammonium assimilation, which limits nitrogen available to symbionts and constrains their growth [56, 57]. The aquaporins AQP3 and AQP5 were also activated following 18 h of heat stress, although their substrate specificity in Aiptasia has not been tested [58]. Immune-related activation targets included GBP1, an interferon-induced GTPase that targets bacteria in vertebrates [59], and PPP3CA, the catalytic subunit of calcineurin, a pathway implicated in symbiosis maintenance in Aiptasia [60]. Although TGF-β signaling in cnidarian symbiosis has mainly been studied for its role in immune suppression [32–34], the range of activation targets is consistent with its pleiotropic function as a cytokine and suggests a broader role in symbiosis than immune regulation alone.

### Putative SMAD3 repression targets in symbiotic animals

Putative repression targets were genes with increased SMAD3 binding and lower expression in symbiotic than aposymbiotic animals within the same contrast. These made up a growing share of targets under heat stress, outnumbering activation targets at both 6 h and 18 h (Fig. 5D). Under control conditions, repression targets were enriched for sterol and ceramide transport and monoamine metabolism, and at 18 h for GPCR signaling, amine metabolism, DNA repair, and chromatin organization (Fig. 7A, B). Although serotonin-like immunoreactivity and putative catecholamine pathway components have been reported in cnidarians [61], phylogenomic analyses indicate that the canonical monoaminergic system, including its receptors, is a bilaterian innovation, and these cnidarian sequences cannot be reliably assigned bilaterian functions [62]. We therefore interpret the genes annotated as HTR1A and HRH2 as cnidarian GPCRs of unknown function, which, together with the repression of ADGRD1 and TACR1, indicates that host GPCR expression differs between symbiotic states.

Other repression targets were associated with sterol trafficking and regulation of TGF-β signaling. The most conserved canonical NPC2 paralog (AIPGENE3167), which Hambleton et al. [63] proposed acts as a housekeeping sterol transporter throughout the host, was a putative SMAD3 repression target. In contrast, the non-canonical NPC2 (AIPGENE22527), which localizes to the symbiosome and binds symbiont-derived sterols [63], was not a SMAD3 target but was upregulated in symbiotic animals in all three contrasts, consistent with previous reports [63, 64] (Additional file 5). SMAD3 may therefore contribute to symbiosis-associated sterol handling by repressing the canonical copy, while the non-canonical copy is regulated through other mechanisms. Repression targets also included the co-receptor CD109 and the SMAD3-degrading kinase GSK3 [65, 66], which both negatively regulate TGF-β signaling in vertebrates.

Repression of these inhibitors raises the possibility of positive feedback on SMAD3 activity. Combined with repression of complement C2, a complement component characterized in Aiptasia [67], and activation of GBP1 and PPP3CA, these results suggest that SMAD3-associated regulation of immune genes in symbiotic animals is bidirectional and context dependent. Beyond signaling and immunity, genes annotated as the rRNA methyltransferase ZCCHC4 and the histone acetyltransferase KAT6A were among the most strongly correlated repression targets. ZCCHC4 was repressed under control conditions and at 18 h, consistent with the enrichment of rRNA methylation in both contrasts (Fig. 7A, B), and KAT6A was repressed at 18 h, suggesting that SMAD3-associated repression under prolonged heat extends to chromatin regulators.

## Conclusions

This study provides, to our knowledge, the first genome-wide map of SMAD3 binding in a symbiotic cnidarian and links it to gene expression across symbiotic states and short-term heat stress. SMAD3 binding in Aiptasia was highly condition-specific and differed primarily by symbiotic state, with nearly all differentially bound sites showing higher binding in symbiotic animals and extended to the most sites after 18 h. As heat stress persisted, SMAD3 targets accounted for a growing share of symbiosis-associated expression differences, suggesting that SMAD3 contributes more to the host response as thermal stress is prolonged.

Integrating binding with expression identified 625 putative SMAD3 target genes, including both activated and repressed genes, supporting a bifunctional, context-dependent role for SMAD3. Activated targets connected TGF-β signaling to the mucociliary surface, nitrogen assimilation, retinoid uptake, and immunity. Repression targets included sterol trafficking genes, cnidarian GPCRs, and negative regulators of TGF-β signaling, pointing to a possible positive feedback loop that sustains SMAD3 activity. Together, these results suggest that TGF-β/SMAD3 signaling in cnidarians extends beyond immune suppression to coordinate nutrient exchange, retinoid signaling, and immune regulation at the symbiotic interface. These putative targets provide a framework for testing how TGF-β signaling supports cnidarian-algal symbiosis and how this regulation responds to the thermal stress that drives coral bleaching.

## Methods

### Aiptasia husbandry

Aiptasia (*Exaiptasia diaphana*) from the CC7 clonal line, symbiotic with *Symbiodinium linucheae*, were maintained in 1 L autoclaved seawater (37 ppt, obtained from the Red Sea) at 25°C under 40 μmol photons m^-2^s^-1^ of light on a 12-h light:12-h dark cycle. Symbiotic and aposymbiotic animals were both kept in transparent containers (Cambro 36CW135) and fed freshly hatched *Artemia salina* nauplii three times weekly with water changes occurring 6–8 h after feeding. Aposymbiotic animals were generated by menthol bleaching [68] and were light acclimated for at least two months prior to sampling.

### Antibody design

A custom rabbit polyclonal antibody was generated by GenScript against an Aiptasia SMAD3-specific peptide (MSGDQQQNMQTDQQC). The full Aiptasia SMAD3 amino acid sequence (Accession no. KXJ28546.1) was obtained from the Aiptasia genome [35] on NCBI. Peptide selection was guided using InterPro v. 98.0 [69] and PredictProtein [70] to identify an exposed amino acid region outside the DNA-binding, protein-binding, or phosphorylation domains. The chosen sequence was predicted to be unique to SMAD3, minimizing potential cross-reactivity with related proteins. Antibody purification and validation were performed by GenScript following the manufacturer’s standard protocols. Western blotting with the SMAD3 antibody detected and confirmed a band at 48 kDa (Additional file 1: Fig. S1A). Immunofluorescence staining detected SMAD3 in the gastrodermis and epidermis in both symbiotic and aposymbiotic anemones (Additional file 1: Fig. S1B).

### Experimental design

To investigate SMAD3 binding and transcriptional responses to acute heat stress, aposymbiotic and symbiotic Aiptasia (0.5–1 cm oral disk diameter) were exposed to control conditions or 6 h or 18 h of heat stress, for six conditions in total (Fig. 2A). Animals were acclimated for two weeks in experimental tubs (Cambro 92CW135) containing 300 mL of autoclaved seawater, with 10–20 animals per tub and three tubs per condition as biological replicates (18 samples total). Control tubs remained at 25°C and were sampled at 6 h (CA, CS), while heat-stressed tubs were moved to a 34°C incubator and sampled at 6 h (HA6, HS6) and 18 h (HA18, HS18) (Fig. 2A). Because controls were sampled only at 6 h, the 18 h comparisons cannot separate heat duration from time-dependent effects. From each tub, 5–10 animals were pooled for ChIP-seq and 1–3 for RNA-seq to obtain sufficient starting material (Additional file 2: Table S1A), then snap-frozen in liquid nitrogen and stored at −80°C.

### Chromatin immunoprecipitation

The chromatin immunoprecipitation protocol was adapted from Dix et al. [71] (Fig. 2B). Animals were removed from −80°C and immediately placed into 10 mL of cross-linking buffer (2 mM disuccinimidyl glutarate, 50 mM EDTA, 25 mM EGTA, 1X DPBS) in a 15 mL conical tube and rotated at 12 rpm for 45 min at room temperature (Stuart SB3 Rotator). Samples were briefly centrifuged, the buffer was decanted, then 10 mL of formaldehyde cross-linking buffer (1% formaldehyde, 50 mM EDTA, 25 mM EGTA, 1X DPBS) was added, and rotated at 12 rpm for 1.5 h at 4°C. Cross-linking was quenched in 10 mL of glycine buffer (0.125 M glycine, 50 mM EDTA, 25 mM EGTA, 1X DPBS) with rotation for 20 min at 4°C. Animals were washed once with cold 1X DPBS and snap-frozen in liquid nitrogen until further processing. Between each buffer incubation, samples were briefly centrifuged and the supernatant was decanted.

Tissue was homogenized in liquid nitrogen with a mortar and pestle, then added to cold lysis buffer (10 mM Tris-HCl, 10 mM NaCl, 50 mM EDTA, 25 mM EGTA, 0.1% Na-Deoxycholate, 0.5% N-Lauroyl sarcosine, 1% SDS, 1X protease inhibitor cocktail). The lysate was then passed through a Wheaton glass tissue grinder and a 25G needle to break up tissue aggregates. Samples were sonicated on ice and chromatin fragment size was confirmed to be 150–500 bp using gel electrophoresis before proceeding (see [71] for the detailed protocol). The salt concentration of the lysate was raised from 10 mM to 150 mM and centrifuged at 20,000 × g for 20 min at 4°C. The resulting chromatin pellets were resuspended in cold immunoprecipitation (IP) buffer (10 mM Tris, 140 mM NaCl, 1 mM EDTA, 0.5 mM EGTA, 1% Triton X-100, 0.1% Na-Deoxycholate, 1X protease inhibitor cocktail). Ten percent of the volume was removed for the input control sample and stored at −20°C. The remaining volume was split for the no-antibody mock IP and SMAD3 IP using 5 µg antibody and rotated at 12 rpm overnight at 4°C.

For immunocomplex recovery, 50 µL of magnetic beads (DynaBeads protein A; ThermoFisher) per IP were first blocked in 0.5% BSA in 1X DPBS by rotating at 12 rpm for 30 min at 4°C, then added to SMAD3 IP samples, and rotated at 12 rpm for 3 h at 4°C. Immunocomplexes were separated on a magnetic rack and alternately washed twice with a low salt buffer (150 mM NaCl, 20 mM Tris-HCl (pH 8), 1% Triton X-100, 2 mM EDTA, 0.1% SDS) followed by a high salt buffer (500 mM NaCl, 20 mM Tris-HCl (pH 8), 1% Triton X-100, 2 mM EDTA, 0.1% SDS). They were washed twice with TE buffer (10 mM Tris, 1 mM EDTA, 50 mM NaCl), resuspended in an elution buffer (50 mM Tris-HCl (pH 8), 10 mM EDTA, 1% SDS), and incubated in a Thermomixer (Eppendorf) at 700 rpm for 15 min at 65°C. The eluate, now separated from the beads, was transferred to a new tube. The corresponding input samples were removed from the freezer, and an equal volume of elution buffer was added. All samples were placed back in the Thermomixer and reverse cross-linked by shaking at 700 rpm overnight at 65°C. RNA and protein were digested, and DNA was extracted using phenol:chloroform:isoamyl alcohol 25:24:1 (Sigma-Aldrich) according to the manufacturer’s instructions. DNA concentrations were measured using Qubit dsDNA broad range and high-sensitivity kits before library preparation.

### RNA extraction

RNA was extracted using TRIzol in combination with an RNeasy Mini kit (Qiagen). Samples were transferred from −80°C to liquid nitrogen and immediately homogenized in TRIzol using a mechanical homogenizer for 3 min. To remove tissue debris, samples were centrifuged at 12,000 × g for 10 min at 4°C. Chloroform was added to the supernatant, vortexed, and centrifuged at 12,000 × g for 15 min at 4°C. Isopropanol was added to the aqueous layer, vortexed, and incubated at room temperature for 15 min. Samples were transferred to an RNeasy spin column and were processed further according to the manufacturer’s instructions (RNeasy Mini kit 74104). RNA concentrations were measured using a Qubit RNA high-sensitivity kit and RNA quality was assessed with an Agilent TapeStation before cDNA library preparation (Additional file 2: Table S1D).

### Library preparation and sequencing

ChIP-seq libraries were prepared using the NEBNext® Ultra™ II DNA Library Prep Kit (Illumina) for the input and SMAD3 samples with starting DNA quantities ranging from 5.1–150 ng. Adapter dilution, PCR enrichment, and fragment size selection were determined based on the kit’s recommendations (Additional file 2: Table S1B). Briefly, adaptor dilution was omitted if the total DNA input was less than 100 ng, otherwise adaptors were diluted 1:10 before adaptor ligation. For samples with less than 50 ng of starting material, fragment size selection was not performed, and 11 PCR cycles were performed to amplify the adaptor-ligated DNA. If there was more than 50 ng of starting material, fragment size selection was performed, and 7 PCR cycles were performed.

RNA-seq poly(A) enriched cDNA libraries were prepared using the NEBNext® Ultra™ II Directional RNA Library Prep Kit (Illumina) with a starting RNA input of 1 µg. ChIP-seq and RNA-seq library concentrations were measured using Qubit dsDNA broad range and high-sensitivity kits. Library quality was assessed using a high sensitivity dsDNA D5000 ScreenTape assay on the Agilent TapeStation. For ChIP-seq, the average insert size was 278 bp. For RNA-seq, the average fragment size was 410 bp. Final ChIP-seq and RNA-seq libraries were pooled in equimolar concentration prior to sequencing, yielding 72 total libraries: 54 ChIP-seq (six conditions × three replicates × three sample types (input, mock, and SMAD3)) and 18 RNA-seq (six conditions × three replicates). Libraries were sequenced as 2 x 150 bp paired-end reads using one 25B lane on the Illumina NovaSeq X platform through BMKGene (Hong Kong). No-antibody mock libraries were used for initial quality checks of ChIP enrichment and were not included in downstream analyses.

### Trimming and alignment

Raw read pairs had a median depth of 49 million (mean 64 million, min 650,000, max 656 million) and were quality-trimmed using TrimGalore v. 0.6.7 [72] with Cutadapt v. 4.9 [73] to remove adapter sequences, low-quality bases, and poly(A) contamination. For ChIP-seq samples, trimmed reads were aligned to a combined reference genome of Aiptasia from the CC7 clonal line (Accession no. GCA_001417965.1; [35]) and *Symbiodinium linucheae* (Accession no. GCA_905221635.1; [74]) using Bowtie2 v. 2.5.4 [75] in local alignment mode (--local -L 16 --score-min L,16,1 --no-unal). The average alignment rate was 88.1% (Additional file 2: Table S1C). Aligned reads with mapping quality scores less than 30 were filtered out using Samtools v. 1.21 [76] to remove multi-mapped reads. Reads mapping to *S. linucheae* were discarded, and the remaining host-aligned reads were deduplicated using Picard v. 3.1.1 [77] with the function MarkDuplicates, yielding a final median depth of 44 million unique, high-quality aligned reads per sample (mean 55 million, min 459,000, max 550 million) (Additional file 2: Table S1C).

### Peak calling

To mitigate the effects of uneven sequencing depth on peak calling, high-quality reads were randomly subsampled to a uniform target depth for each sample using Samtools (samtools view -s). Target depths were set at 40 and 25 million reads for input and SMAD3 samples, respectively to maximize the number of samples with even depth and minimize read loss. This normalization step reduced the influence of uneven sequencing depth on peak detection. Samples already below these thresholds were retained at their original depth (Additional file 1: Fig. S2A; Additional file 2: Table S1C).

Regions of SMAD3 enrichment (peaks) were called on the subsampled BAM files using MACS3 v. 3.0.2 [78] with matched input controls as background. Peaks were called in paired-end mode, using fragment sizes from read pairs, with an effective genome size of 2.06 × 10^8^ and a p-value threshold of 0.05. The fraction of reads in peaks (FRiP) was calculated as the number of reads in peaks divided by the total number of mapped reads and was used to assess ChIP enrichment quality. This resulted in an average of 40,752 called peaks per sample (Additional file 1: Fig. S2B) and an average FRiP of 14.9% (Additional file 1: Fig. S2C). The ENCODE benchmark for an acceptable FRiP (>1%) was established using peaks called under MACS’s default q < 0.05 threshold in mammalian genomes [79]; because FRiP scales with the leniency of the peak-calling threshold itself (FRiP and peak number were positively correlated across samples, R² = 0.705), our values are not directly comparable to this guideline.

Rather than relying on per-sample enrichment strength as a quality filter, we used a permissive initial threshold to maximize sensitivity and instead required peaks to be reproducible across biological replicates. MACS3-called peaks were then processed with Multiple Sample Peak Calling (MSPC) v. 1.15.0 [80, 81] to evaluate reproducibility across biological replicates and to remove false-positive peaks. MSPC classifies each peak as stringent or weak based on the MACS3-reported p-value. Peaks with a p-value below the stringency threshold of 1 × 10^-5^ were classified as stringent, while those between 1 × 10^-5^ and the weak threshold of 1 × 10^-2^ were classified as weak. Weak peaks that would otherwise be discarded can be rescued if they overlap with peaks in additional replicates, and joint significance is assessed by combining p-values across replicates using Fisher’s method. MSPC was run separately for each condition (symbiotic state × treatment) with replicates treated as biological replicates. Peaks passing the reproducibility filter in each replicate were called confirmed peaks. Consensus peaks were then generated by merging confirmed peaks across all replicates within a condition into a single non-redundant peak set. With c = 2, a peak required support from at least two of three replicates to be confirmed and resulted in an average of 3,020 peaks per replicate (Additional file 1: Fig. S2D) and 3,504 consensus peaks per condition (Fig. 3A). MSPC was also run with c = 3, requiring support from all three replicates, to generate a high-confidence consensus peak set for overlap visualization (Additional file 1: Fig. S3).

To assess the overlap of SMAD3 binding sites across conditions, the consensus peaks for each condition were merged by genomic coordinates into a unified set of non-redundant loci, with 17,598 total peaks for c = 2 and 3,309 for c = 3. A binary presence/absence matrix was constructed by intersecting each condition’s consensus peaks against the merged loci, and the resulting overlap patterns were visualized as UpSet plots with ComplexUpset v. 1.3.3 [82].

### Genomic annotation of peaks

Genomic annotation of SMAD3 binding sites was performed using ChIPseeker v. 1.46.1 [83]. A transcript database was constructed from the Aiptasia genome annotation from the NCBI RefSeq GTF (GCF_001417965.1) and MSPC consensus peaks from all six conditions were annotated to genomic features. Peaks were classified into genomic features including 5’ UTR, 3’ UTR, exon, intron, downstream (≤ 300 bp), distal intergenic regions, and promoter regions, defined as being within ±3 kb of the transcription start site (TSS). The distribution of peaks across genomic features was visualized across all conditions and as a combined pie chart with default parameters. Genomic features representing less than 0.05% of peaks were omitted from the pie chart.

### Differential binding analysis

Differential binding analysis was performed using DiffBind v. 3.12.0 [84, 85], which uses DESeq2 v. 1.50.1 [86] for statistical modeling. MSPC-confirmed peaks per sample (c = 2) were used as input peak locations. Each peak was re-centered on its point of highest enrichment and extended 200 bp in each direction (summits = 200), creating uniform 401-bp windows for read counting. Reads overlapping each window were counted from the original, non-subsampled BAM files to maximize statistical power, producing a count matrix of 17,569 binding sites across all 18 samples. Library sizes were estimated from reads falling outside peak regions (library = DBA_LIBSIZE_BACKGROUND) to prevent differences in enrichment efficiency from biasing normalization. Read counts were modeled using a negative binomial distribution in DESeq2 with an interaction model (design = ∼ symbiotic state × treatment) to test the individual and combined effects of symbiotic state (aposymbiotic, symbiotic) and heat stress treatment (control, 6 h, 18 h) on SMAD3 binding intensity. An additive model (design = ∼ symbiotic state + treatment) was also fit to estimate the overall effects of symbiotic state and heat treatment across all samples. Pairwise contrasts were extracted to assess the effect of symbiotic state at each heat stress timepoint (CS vs CA, HS6 vs HA6, HS18 vs HA18), the effect of heat stress within each symbiotic state (HS6 vs CS, HS18 vs CS, HA6 vs CA, HA18 vs CA, HS6 vs HS18, HA6 vs HA18), and the interaction of heat treatment and symbiotic state. Binding sites were considered significantly differentially bound at an adjusted p-value < 0.05 (Benjamini-Hochberg correction).

Variance-stabilized count data were generated in DESeq2 with blind = TRUE and a local fit. Principal component analysis (PCA) was performed to visualize sample clustering by symbiotic state and treatment. To statistically assess clustering, PERMANOVA was performed on Euclidean distance matrices of variance-stabilized counts using the adonis2 function in vegan v. 2.7.5 [87] with 999 permutations, with symbiotic state, treatment, and their interaction as model terms. Homogeneity of multivariate dispersion was verified using the betadisper function.

### Gene expression analysis

For RNA-seq samples, paired reads were pseudo-aligned to a kallisto index built from a combined reference transcriptome of Aiptasia [35] and *S. linucheae* [74]. Transcript-level abundances were estimated using kallisto v. 0.46.2 [88] with 100 bootstrap iterations. The average percent of reads pseudoaligned to the combined transcriptome was 61.8% (Additional file 2: Table S1D). Symbiont-mapped reads were removed, leaving an average of 19.3 ± 6.4 million (mean ± SD) estimated host read counts per sample (Additional file 2: Table S1D).

Transcript abundance estimates were imported into R using tximport v. 1.38.1 [89]. Transcripts with fewer than ten total counts across all samples were removed, resulting in a total of 27,139 transcripts. Variance-stabilized count data were generated using DESeq2 [86] and PCA was performed with default settings and visualized using ggplot2 v. 4.0.1 [90]. Effects of symbiotic state and heat stress treatment on global expression profiles were assessed using the same PERMANOVA framework described for differential binding analysis.

Differential gene expression was analyzed with DESeq2 v. 1.50.1 [86] and modeled as a function of symbiotic state (aposymbiotic, symbiotic), treatment (control, 6 h, 18 h), and their interaction using the model: design = ∼ symbiotic state × treatment. Pairwise contrasts were extracted to assess the effect of symbiotic state within each treatment level (CS vs CA, HS6 vs HA6, and HS18 vs HA18), the effect of heat stress within each symbiotic state, and interaction terms testing whether the transcriptional response to heat stress differed between symbiotic states. Transcripts were considered differentially expressed at an FDR-adjusted p-value < 0.05 using a Benjamini-Hochberg correction with a log2 fold change threshold of ±1. Full differential expression results for all contrasts are provided in Additional file 5.

### Peak to gene mapping and correlation

Gene coordinates and strand information were extracted from the NCBI RefSeq annotation of the Aiptasia genome (GCF_001417965.1; 24,862 annotated genes), and promoter-proximal regions were defined as ±3 kb from each TSS using GenomicRanges v.1.62.0 [91]. RNA-seq gene models (GenBank; GCA_001417965.1) [35] were matched to RefSeq genes by maximum same-strand coordinate overlap. Each differentially bound SMAD3 peak was reduced to its midpoint and assigned to every gene whose promoter-proximal region contained it, so a single peak could be assigned to more than one gene. Genes with a differentially bound peak in their promoter-proximal region that were also differentially expressed in the corresponding RNA-seq contrast were considered putative SMAD3 transcriptional targets. These target genes were then split by the direction of their RNA-seq fold change into activation targets (increased SMAD3 binding with upregulated expression in symbiotic animals) and repression targets (increased binding with downregulated expression) (Fig. 5B). To assess whether identifier matching affected the target set, peak-to-gene assignment was repeated using TSSs from the AIPGENE GenBank gene models, allowing differentially bound peaks to be intersected directly with all DEGs without identifier matching. Using this set of DEGs containing differentially bound SMAD3 peaks in their promoter-proximal region, we examined the relationship between SMAD3 promoter binding and target gene expression at the level of individual genes. Pearson’s correlation coefficient (r) was calculated between variance-stabilized SMAD3 binding and expression across the 18 paired samples as a descriptive measure of binding–expression concordance. Because samples were grouped by condition and targets were selected for differences between symbiotic states, r primarily reflects symbiotic-state separation and was used descriptively rather than as an independent test. For a select set of genes with higher Pearson correlation coefficients, scatter plots were generated with ggplot2 [90], colored by condition and grouped by functional category. For each target gene, the differentially bound promoter-proximal peak with the smallest peak-midpoint-to-TSS distance was selected to represent its binding signal. Per-gene binding and expression values were taken from non-blind variance-stabilized transformed count matrices. Candidate genes for scatter plots were selected from Additional file 7 to represent functional categories among targets with higher correlation coefficients.

Binding and Expression Target Analysis (BETA) v. 2.1.1 was used to predict regulatory gene targets and infer transcription factor function by integrating ChIP-seq and RNA-seq data [38]. The full consensus SMAD3 peak set (17,569 peaks) was supplied as the binding input, scored by mean library-size-normalized signal across all ChIP-seq samples. Differential gene expression results for each of three symbiotic-state contrasts were obtained from the DESeq2 interaction model described above. BETA was run separately for each contrast against the same SMAD3 peak set, using a search window of ±3 kb from the TSS and an FDR threshold of 0.05 for defining differentially expressed genes. All called peaks were included in the regulatory potential calculation. BETA calculated a distance-weighted regulatory potential score per gene from all peaks within the search window and used a one-tailed Kolmogorov-Smirnov (KS) test to assess whether up- or downregulated genes showed significantly higher regulatory potential than non-differentially-expressed background genes, applying a significance threshold of p < 0.05 to call transcriptional activation or repression function. Regulatory potential scores were calculated for each gene as the sum of all SMAD3 peaks within the search window, weighted by distance from the TSS so that closer peaks contribute more. Genes were then ranked by regulatory potential and by differential expression, and the two ranks were combined into a rank product, with lower values indicating stronger combined support. BETA predictions were then cross-referenced with the full set of putatively SMAD3-targeted genes from the three symbiotic-state contrasts.

### Gene ontology (GO) enrichment analysis

Gene ontology (GO) enrichment analysis was performed using topGO v. 2.62.0 [92] with the weight01 algorithm and Fisher’s exact test for biological process terms. GO terms with less than 5 annotated genes were omitted from the analysis and the top 20 most significant GO terms were reported. Aiptasia GO annotations were obtained from the NCBI RefSeq annotation (GCF_001417965.1), yielding 15,973 GO-annotated genes out of 24,862 total genes. DEGs containing a SMAD3 differentially bound peak in its promoter-proximal region were split into two categories: increased binding with upregulated expression or increased binding with downregulated expression. Directional analyses with gene ontology were only performed in contrasts with more than 50 significant genes, which included only CS vs CA and HS18 vs HA18 (Additional file 7). For all GO analyses, the background was restricted to the 12,893 genes that were detectable in the RNA-seq experiment and carried GO annotations, representing the universe of expressed, annotated genes testable in this dataset. For the RNA-seq analysis alone, upregulated and downregulated DEGs were tested separately, and the top 30 GO terms were reported.

## Additional Files

**Additional file 1: Supplementary methods and results. Figure S1.** Validation of custom Aiptasia SMAD3 antibody. **Figure S2.** Peak calling summary information. **Figure S3.** Condition specificity of high-confidence SMAD3 peaks. **Figure S4.** Differential binding contrasts within each symbiotic state. **Figure S5.** Transcriptome-wide gene expression differences between symbiotic states and in response to heat stress. **Figure S6.** GO terms enriched among genes differentially expressed between symbiotic and aposymbiotic animals under control conditions. **Figure S7.** GO term enrichment for genes differentially expressed between symbiotic and aposymbiotic animals under heat stress. **Figure S8.** GO term enrichment for heat-responsive genes in aposymbiotic animals. **Figure S9.** GO term enrichment for heat-responsive genes in symbiotic animals. **Figure S10.** GO term enrichment for genes with a significant symbiotic state × treatment interaction. **Figure S11.** BETA activation and repression function prediction for SMAD3.

**Additional file 2: Table S1.** Sample metadata and sequencing library summaries for ChIP-seq and RNA-seq. (**A**) Sample breakdown for sampling and experimental design. (**B**) ChIP-seq library preparation. (**C**) ChIP-seq library read count summary from Bowtie2 (BT2) alignment. (**D**) RNA-seq library preparation and kallisto pseudoalignment estimated read count summary. (XLS 22 kb)

**Additional file 3: Table S2.** Summary of differential expression and SMAD3 binding-expression integration. (**A**) Summary of up- and down-regulated differentially expressed genes (DEGs) in each condition contrast. (**B**) Summary of SMAD3 differential binding and differential gene expression integration for contrasts between symbiotic states. (**C**) Comparison of putative SMAD3 targets identified using RefSeq (LOC) and GenBank (AIPGENE) gene models. (XLS 14 kb)

**Additional file 4:** Known transcription factor motifs enriched in SMAD3 consensus peaks per condition. (XLS 15 kb)

**Additional file 5:** Differentially expressed genes for all contrasts. (XLS 5.1 mb)

**Additional file 6:** RNA-seq topGO results for all contrasts. (TSV 140 kb)

**Additional file 7:** Putative SMAD3-targeted genes. (XLS 81 kb)

## Abbreviations

CA: control aposymbiotic
CS: control symbiotic
HS6: heat stress 6 h symbiotic
HS18: heat stress 18 h symbiotic
HA6: heat stress 6 h aposymbiotic
HA18: heat stress 18 h aposymbiotic
TGF-β: Transforming growth factor beta
SMAD3: Mothers against decapentaplegic homolog 3
IP: immunoprecipitation
ChIP-seq: Chromatin immunoprecipitation sequencing
DEGs: Differentially expressed genes
TSS: transcription start site
GO: Gene ontology
EDTA: ethylenediaminetetraacetic acid
EGTA: ethylene glycol-bis(β-aminoethyl ether)-N,N,N’,N’-tetraacetic acid
SDS: sodium dodecyl sulfate
DPBS: Dulbecco’s phosphate-buffered saline
NaCl: sodium chloride
Tris-HCl: Tris hydrochloride
UTR: untranslated region
BAM: binary alignment map
PERMANOVA: permutational multivariate analysis of variance
PCA: principal component analysis
ROS: Reactive oxygen species

## Declarations

### Ethics approval and consent to participate

Not applicable

### Consent for publication

Not applicable

### Availability of data and materials

The sequencing data reported in this study has been deposited to NCBI BioProject accession number PRJNA1544129 (https://www.ncbi.nlm.nih.gov/bioproject/PRJNA1544129/). All data analysis scripts are available on GitHub at https://github.com/burleigo/Aiptasia_SMAD3_ChIP-seq.

### Authors’ contributions

O.L.B., M.F.D., G.C., M.A., and V.M.W. conceived the project. O.L.B., M.F.D., and G.C. developed the methodology. O.L.B. performed the experiments. V.M.W. and M.A. provided resources and supervised the project. O.L.B. analyzed and visualized the data with supervision from M.R., M.F.D., G.C., and M.A. O.L.B. prepared the original draft of the manuscript. All authors reviewed, edited, and approved the final manuscript.

## Acknowledgements

We would like to thank the KAUST Bioscience Core Lab facilities and Anne-Marie Girard at the Confocal Microscopy Facility of the Center for Quantitative Life Sciences at Oregon State University.

