## Additional file 1 for "ChIP-seq analysis of SMAD3 target genes in a sea anemone reveals symbiosis-dependent regulation through TGF-β signaling"

**Supplemental Methods**

**Western blots**

Western blots were performed to confirm antigen specificity of the SMAD3 antibody using methods modified from Sawiccy et al. [1]. Briefly, whole symbiotic and aposymbiotic anemones were frozen at -80°C and homogenized in cold lysis buffer (100 mM Tris, 100 mM NaCl, 10 mM EDTA, 1X protease inhibitor cocktail, 1% Tween-20) and centrifuged at 14,000 x g for 15 min at 4°C. The supernatant was transferred to a new tube, and soluble protein was quantified using a Bradford protein assay (ThermoFisher). Forty µg of protein lysate was mixed with 2X Laemmli sample buffer (Bio-Rad) and 5% beta-mercaptoethanol and heated at 95°C for 10 min. Proteins were resolved via SDS-PAGE in a 4-20% gradient gel (Mini Protean TGX stain-free precast gel; Bio-Rad) with a BLUEstain 11-245 kDa protein ladder (GoldBio). The gel was run at 160 V for 75 min at 4°C. Proteins were electrophoretically transferred onto a nitrocellulose membrane in a methanol transfer buffer at 75 V for 100 min at 4°C. The membrane was stained with Ponceau to check for transfer quality and then rinsed with nanopure water and Tris-buffered saline with 0.1% Tween (TBST) to remove the stain. The membrane was blocked in a blocking solution (5% w/v non-fat milk in TBST) with gentle rocking for 1 hour at RT then incubated with anti-SMAD3 primary antibody diluted in blocking solution at a 1:1000 concentration overnight at 4°C. The membrane was washed with TBST three times for 5 min each and incubated with a goat anti-Rabbit IgG-HRP secondary antibody (Santa Cruz) diluted in blocking solution at a 1:5000 concentration for 1 hour at RT and then washed again with TBST. Protein bands were visualized by chemiluminescence using Clarity Western ECL substrate (Bio-Rad) and imaged on an Azure imager.

**Immunolocalization of SMAD3**

Immunofluorescence staining was performed on cryosectioned tissue to look for the presence of SMAD3 localization in the cytoplasm and nucleus. Aiptasia were relaxed in 0.5 M MgCl2, fixed in 4% paraformaldehyde for 2 hours, rinsed in PBS, and cryoprotected in a series of 10%, 20% and 30% sucrose in PBS overnight. Samples were embedded in OCT and cryosectioned at 10 µm. Tissue sections were blocked for 1 h (5% normal goat serum, 0.2% Triton X-100, PBS) and incubated overnight at 4°C in the Aiptasia-specific SMAD3 primary antibody at 1:1000 (GenScript). Sections were washed 3X with PBS-Tween (0.05%), incubated in a goat anti-rabbit IgG Alexa Fluor-488 secondary antibody at 1:500 (Invitrogen) for 2 h, and washed 2X with PBS-Tween. Hoechst 33342 (ThermoFisher) was applied at a final concentration of 1 µg/mL for 5 min followed by two washes in PBS-Tween and one wash in PBS. Cryosections were mounted in Prolong gold antifade reagent (Invitrogen) and imaged on a Leica Stellaris confocal microscope with an HC PL APO 63×/1.40 NA oil immersion objective using sequential acquisition. Hoechst 33342 and chlorophyll autofluorescence were excited at 405 nm and collected at 425–495 nm and 626–799 nm, respectively, and Alexa Fluor 488 was excited at 498 nm using a white light laser and collected at 503–626 nm. Z-stacks of 15 optical sections were acquired at 1 µm intervals with a 1 Airy unit pinhole, a pixel size of 0.18 µm, 600 Hz bidirectional scanning, and 4X line averaging.

**SMAD3 motif enrichment**

Motif enrichment within SMAD3 binding sites was assessed with HOMER v. 4.1 findMotifsGenome.pl. For each of the six conditions, MSPC consensus peaks were analyzed against the Aiptasia genome [2]. Searches used the full width of each consensus peak, with de novo motif lengths of 6, 8, 10, 12, 14, 16, and 20 bp. HOMER selected background sequences from random genomic positions, sized to the average width of the target peaks for each condition and matched to their GC content, with additional auto-normalization for short oligonucleotide composition. Enrichment of known motifs from the full HOMER known motif collection was tested with a cumulative binomial test, and motifs with a Benjamini-Hochberg q-value below 0.05 were considered significantly enriched.

**Supplemental Results**

**Heat stress elicits a rapid transcriptional response peaking at 6 hours**

RNA-seq was used to characterize transcriptomic differences between symbiotic states and in response to heat stress. PCA of blind variance-stabilized expression separated samples by symbiotic state along PC1, which explained 70% of the variance, and by heat treatment along PC2, which explained 12% (Fig. S5A). PERMANOVA confirmed that symbiotic state explained the largest proportion of transcriptomic variance (R² = 0.45, p = 0.001), followed by heat treatment (R² = 0.20, p = 0.004), with no significant interaction (R² = 0.076, p = 0.124). Multivariate dispersion was homogenous for both factors (betadisper, p > 0.18). Differential expression was assessed using the same interaction model as for differential binding (design = ~ Symbiotic state × Treatment). Differential expression was assessed with the same interaction model used for differential binding (design = ~ Symbiotic state × Treatment). Expression divergence between symbiotic states was greatest at 6 h of heat stress, with 12,319 DEGs in HS6 vs HA6 compared with 7,876 in CS vs CA and 5,535 in HS18 vs HA18 (Fig. S5B–D, Table S2A, File S1). This pattern contrasts with differential SMAD3 binding between symbiotic states, which was lowest at 6 h and highest at 18 h (Fig. 6C). Within each symbiotic state, the transcriptional response to heat was likewise largest at 6 h and attenuated by 18 h, with 7,019 and 2,303 DEGs in aposymbiotic animals and 5,180 and 3,062 DEGs in symbiotic animals at 6 h and 18 h, respectively (Fig. S5E–H). The symbiotic state × treatment interaction terms, which identify genes whose heat response differs between symbiotic states, yielded 1,664 DEGs at 6 h and 146 at 18 h (Table S2A).

**GO enrichment reveals a heat-induced proteostasis response shared across symbiotic states and a state-specific immune response**

DEGs from each contrast were tested for GO enrichment against a shared background of 12,893 expressed, GO-annotated genes using topGO [3], with upregulated and downregulated genes tested separately to resolve the direction of enrichment (see File S2 for all contrasts and gene ontologies tested). Several processes distinguished symbiotic from aposymbiotic animals regardless of temperature. G protein-coupled receptor signaling pathway (GO:0007186) was consistently and exclusively downregulated in symbiotic animals in CS vs CA (p = 2.1e-10), HS6 vs HA6 (p = 9.0e-20), and HS18 vs HA18 (p = 4.5e-21), while protein polyubiquitination (GO:0000209) was upregulated in all three contrasts (p = 0.0067, 2.6e-6, and 0.0026, respectively; Figs. S6, S7). Cilium movement and related ciliary terms were also upregulated in symbiotic animals under control conditions (Fig. S6A), and vitamin A import into cell was upregulated at 18 h (Fig. S7C). Proteolysis (GO:0006508) and DNA integration (GO:0015074) showed the strongest overall enrichment but were significant among both upregulated and downregulated genes in all three contrasts, consistent with broad GO terms rather than a directionally coherent signature (Figs. S6, S7).

Within each symbiotic state, a core proteostasis response comprising protein folding, protein refolding, cellular response to unfolded protein, and response to endoplasmic reticulum stress was upregulated in all four heat stress contrasts (weight01 Fisher p < 0.05 in each contrast; Figs. S8, S9). Other heat-responsive processes differed between symbiotic states. Regulation of canonical NF-kappaB signaling was upregulated in the HA6 vs CA contrast (p = 0.031) and HA18 vs CA contrast (p = 0.00012), and tumor necrosis factor-mediated signaling was additionally upregulated in the HA18 vs CA contrast (p = 0.0027), but neither was significant in the corresponding symbiotic heat contrasts (Figs. S8, S9). G protein-coupled receptor signaling pathway was upregulated in HA6 vs CA (p = 9.5e-11; Fig. S8A), in contrast to its consistent downregulation in symbiotic relative to aposymbiotic animals (Fig. S6B, S7B, S7D). Ubiquitin-dependent protein catabolic process was upregulated at 6 h in both symbiotic states but was more strongly enriched in symbiotic animals (p = 2.0e-9 in HS6 vs CS, p = 0.0022 in HA6 vs CA; Figs. S8A, S9A).

To identify genes whose response to heat differed between symbiotic states, we examined the symbiotic state × treatment interaction, which compares the heat response in symbiotic animals with that in aposymbiotic animals at each time point. Positive interaction coefficients indicate genes whose expression was relatively higher with heat stress in symbiotic than in aposymbiotic animals, and negative coefficients indicate the reverse. At 6 h, genes with positive coefficients were enriched for innate immune response, cytokine-mediated signaling, inflammatory response, pattern recognition receptor signaling, and positive regulation of canonical NF-kappaB signaling, while genes with negative coefficients were enriched for DNA repair processes, including interstrand cross-link repair and double-strand break repair (Fig. S10A, B). The 18 h interaction yielded far fewer DEGs, and its enrichments were dominated by lipid and amino acid metabolic processes (Fig. S10C, D).

**Supplementary Figures**

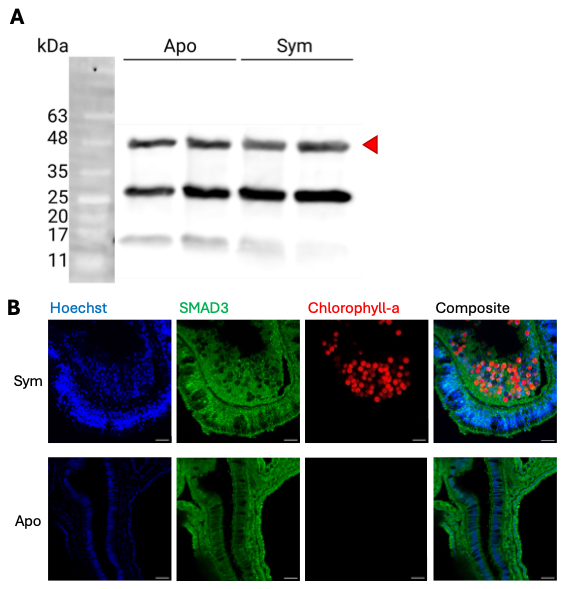

**Figure S1. Validation of custom Aiptasia SMAD3 antibody.**

(**A**) Western blot using aposymbiotic and symbiotic whole-animal lysate (n=2). SMAD3 antibody stains a band at predicted size (~48 kDa; red arrow). Additional bands at ~28-30 kDa and ~15-16 kDa are likely due to proteolytic cleavage of full-length protein. Sym = symbiotic, apo = aposymbiotic. (**B**) Representative maximum projection confocal images of 10-µm sectioned symbiotic and aposymbiotic Aiptasia tissue stained with SMAD3 antibody. Scale bar: 20 µm.

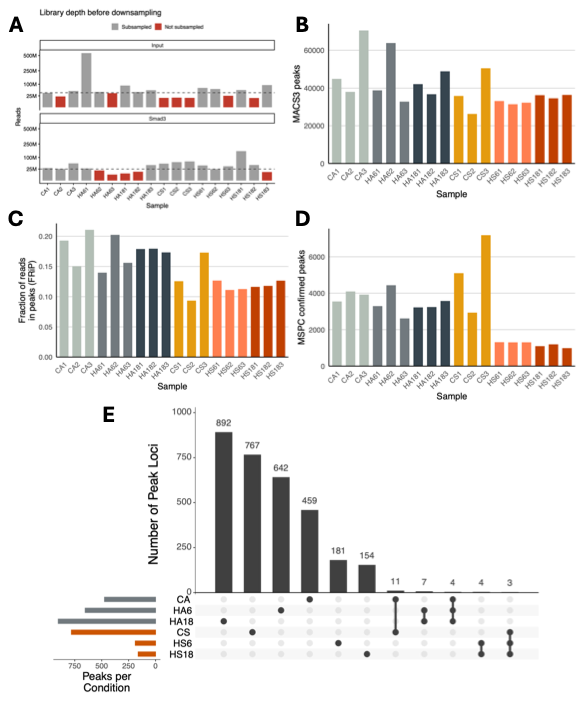

**Figure S2. Peak calling summary information.**

(**A**) Library depth of deduplicated, host-aligned Smad3 and input ChIP-seq BAM files before subsampling. Samples in grey were subsampled to the target depth indicated with the dashed line (Input: 40 million; Smad3: 25 million). Samples in red were not subsampled, and their full depth was used for MACS3 peak calling. (**B**) Number of peaks called by MACS3 after subsampling BAM files. (**C**) Fraction of reads in peaks (FRiP) per sample, calculated as the proportion of mapped reads falling within called peaks. (**D**) Number of MSPC-confirmed peaks per sample, representing peaks that passed reproducibility filtering in at least two of three biological replicates (c = 2). Aposymbiotic samples shown in grey hues, symbiotic samples in orange hues.

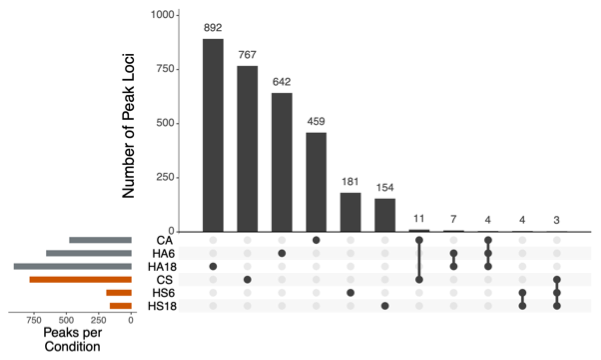

**Figure S3. Condition specificity of high-confidence SMAD3 peaks.**

UpSet plot of MSPC consensus peaks recovered in all three biological replicates of each condition (c = 3), a stricter threshold than the c = 2 consensus set used for downstream analyses. Left bars show the total number of consensus peaks per condition. Top bars show the number of peak loci in each intersection, indicated by connected dots below. Most peaks were unique to a single condition, and only a small number were shared among conditions. Selected intersections between conditions of interest are shown. Grey, aposymbiotic; orange, symbiotic.

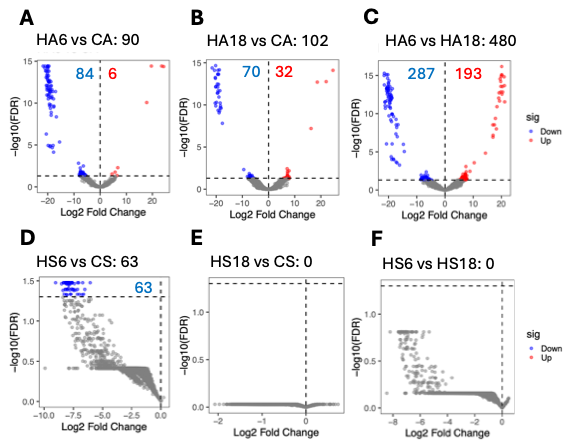
**Figure S4. Differential binding contrasts within each symbiotic state.**

Volcano plots of differential Smad3 binding for within each symbiotic state from the interaction model (design = Symbiotic state x Treatment). Differential Smad3 binding between contrasts in aposymbiotic animals: HA6 vs CA (**A**), HA18 vs CA (**B**), and HA6 vs HA18 (**C**). Differential Smad3 binding between contrasts in symbiotic animals: HS6 vs CS (**D**), HS18 vs CS (**E**), and HS6 vs HS18 (**F**). X-axis, log2 fold change (first condition relative to second condition); y-axis, −log10(FDR). Red, increased binding in first condition; blue, decreased binding in first condition relative to the second condition; grey, not significant (NS). Numbers indicate the count of significant peaks per direction. The dashed horizontal line indicates the significance threshold FDR = 0.05. The total number of significant differentially bound peaks is listed to the right of the contrast name.

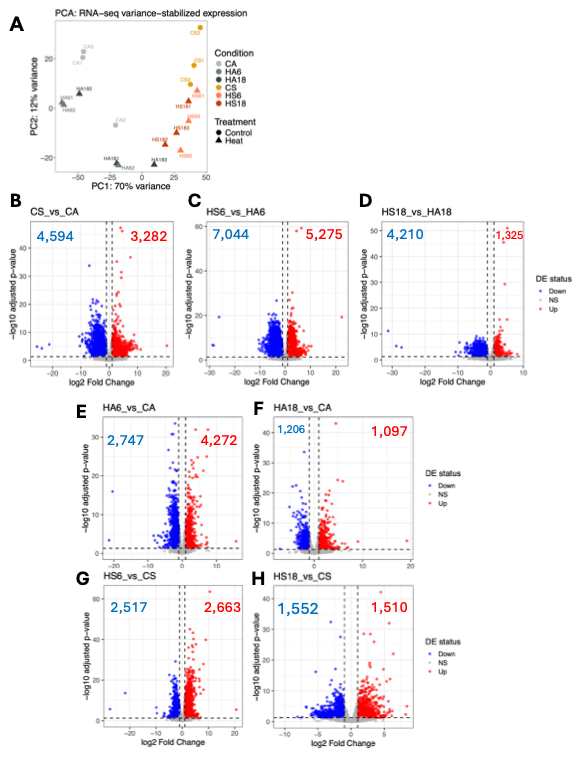
**Figure S5. Differential gene expression.**

(**A**) PCA of RNA-seq variance-stabilized expression (PC1, 70%; PC2, 12% of variance). Aposymbiotic in grey (CA, HA6, HA18), symbiotic in orange (CS, HS6, HS18). (**B-D**) Volcano plots between symbiotic states for control, 6h heat stress, and 18h heat stress with a log2 fold change threshold of ±1 and adjusted p-value threshold of 0.05. Upregulated genes are shown in red; downregulated genes shown in blue; grey is not significant. (**E-H**) Volcano plots within each symbiotic state of heat stress compared to control. Top row is aposymbiotic and bottom row is symbiotic. Left is 6h, right is 18h vs control.

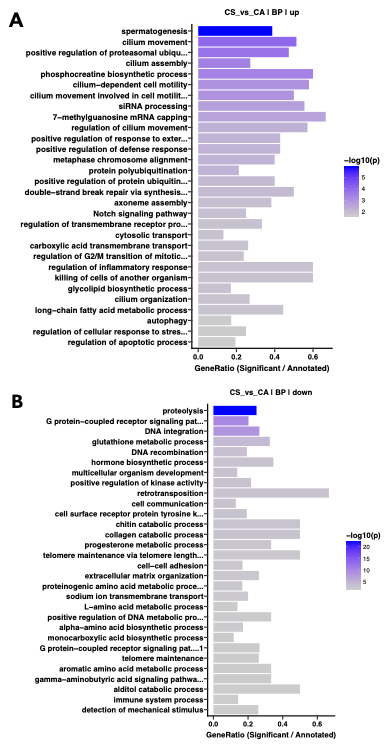

**Figure S6. GO terms enriched among genes differentially expressed between symbiotic and aposymbiotic animals under control conditions.**

Biological process GO terms enriched using topGO among genes upregulated (**A**) or downregulated (**B**) in symbiotic relative to aposymbiotic animals under control conditions (CS vs CA). All expressed genes with GO annotations served as the background. The 30 most significant terms are shown for each direction. Bar color indicates significance (−log10 weight01 Fisher p-value). The x-axis shows the gene ratio (significant genes/annotated genes per GO term). Long term names are truncated.

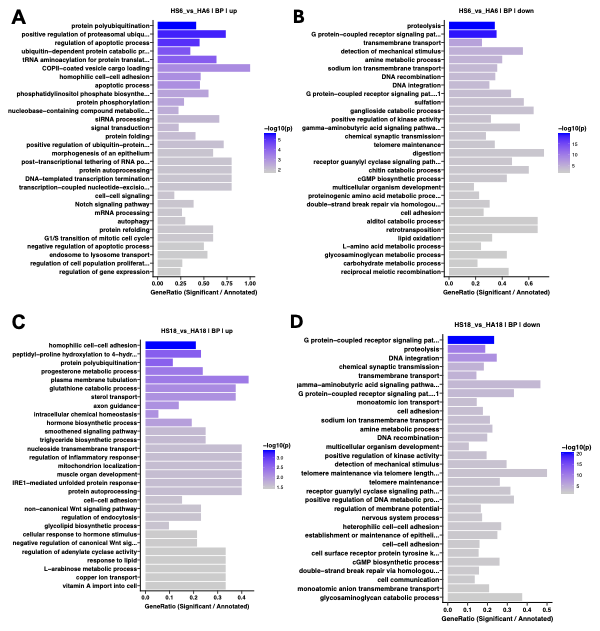
**Figure S7. GO terms enriched among genes differentially expressed between symbiotic and aposymbiotic animals under heat stress.**

Biological process GO terms enriched using topGO among genes upregulated (**A**,**C**) or downregulated (**B**,**D**) in symbiotic relative to aposymbiotic animals after 6 h (**A–B**; HS6 vs HA6) and 18 h (**C–D**; HS18 vs HA18) of heat stress. All expressed genes with GO annotations served as the background. The 30 most significant terms are shown for each direction. Bar color indicates significance (−log10 weight01 Fisher p-value). The x-axis shows the gene ratio (significant genes/annotated genes per GO term). Long term names are truncated.

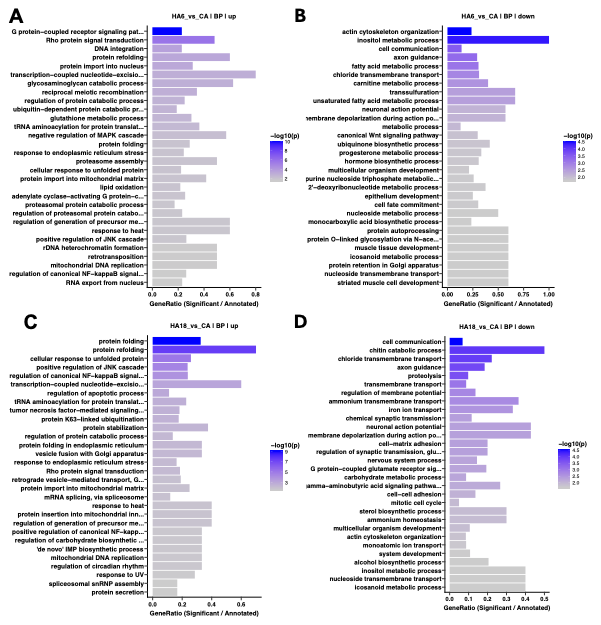

**Figure S8. GO terms enriched among heat-responsive genes in aposymbiotic animals.**

Biological process GO terms enriched using topGO among genes upregulated (**A**,**C**) or downregulated (**B**,**D**) in aposymbiotic animals after 6 h (**A–B**; HA6 vs CA) and 18 h (**C-D**; HA18 vs CA) of heat stress relative to aposymbiotic controls. All expressed genes with GO annotations served as the background. The 30 most significant terms are shown for each direction. Bar color indicates significance (−log10 weight01 Fisher p-value). The x-axis shows the gene ratio (significant genes/annotated genes per GO term). Long term names are truncated.

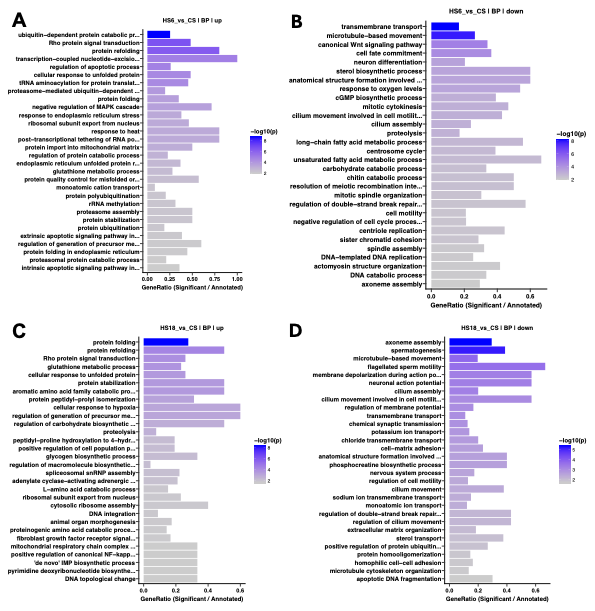

**Figure S9. GO terms enriched among heat-responsive genes in symbiotic animals.**

Biological process GO terms enriched using topGO among genes upregulated (**A**,**C**) or downregulated (**B**,**D**) in symbiotic animals after 6 h (**A–B**; HS6 vs CS) and 18 h (**C-D**; HS18 vs CS) of heat stress relative to symbiotic controls. All expressed genes with GO annotations served as the background. The 30 most significant terms are shown for each direction. Bar color indicates significance (−log10 weight01 Fisher p-value). The x-axis shows the gene ratio (significant genes/annotated genes per GO term). Long term names are truncated.

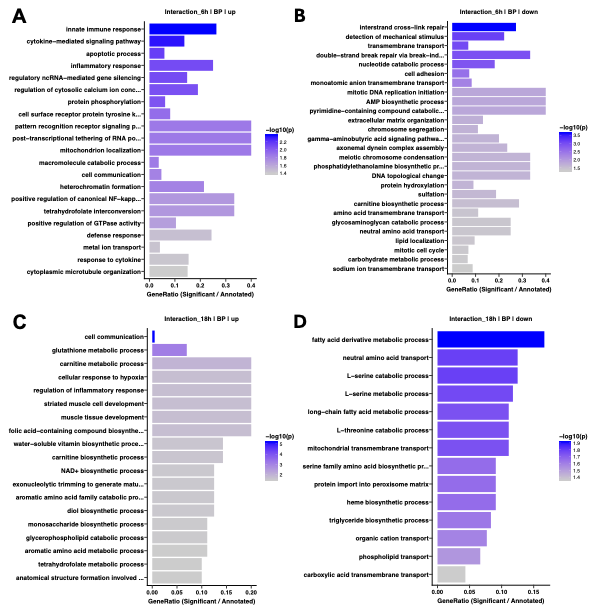

**Figure S10. GO terms enriched among genes with a significant symbiotic state × heat treatment interaction.**

Biological process GO terms enriched using topGO among genes whose transcriptional response to heat differed between symbiotic and aposymbiotic animals, identified by the Symbiotic state × Treatment interaction term in the RNA-seq model. Panels show 6 h (**A–B**; 1,664 DEGs) and 18 h (**C–D**; 146 DEGs). Positive interaction coefficients (up) (**A, C**) indicate genes whose expression was relatively higher after heat stress in symbiotic than in aposymbiotic animals; negative coefficients (down) (**B, D**) indicate the reverse. All expressed genes with GO annotations served as the background. Up to 30 of the most significant terms are shown for each direction. Bar color indicates significance (−log10 weight01 Fisher p-value). The x-axis shows the gene ratio (significant genes/annotated genes per GO term). Long term names are truncated.

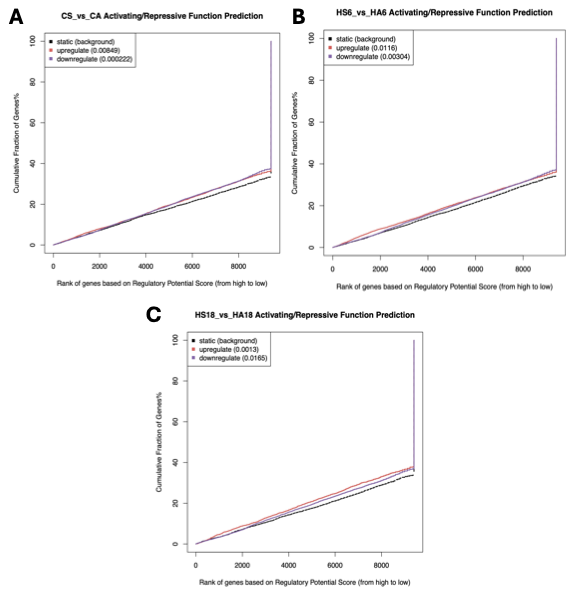

**Figure S11. BETA activation and repression function prediction for SMAD3.**

Cumulative distribution plots for the CS vs CA (**A**), HS6 vs HA6 (**B**), and HS18 vs HA18 (**C**) contrasts. Upregulated (activation-associated) genes are shown in red, downregulated (repression-associated) genes in purple, and non-differentially-expressed background genes in black. A colored curve above the background indicates that the gene set is functionally enriched for SMAD3 binding. P-values (one-tailed KS test vs. background) are shown in parentheses in each panel legend.
